# Pathogenic *Leptospira* are widespread in the urban wildlife of southern California

**DOI:** 10.1101/2023.03.13.531784

**Authors:** Sarah K. Helman, Amanda F.N. Tokuyama, Riley O. Mummah, Mason W. Gamble, Celine E. Snedden, Benny Borremans, Ana C.R. Gomez, Caitlin Cox, Julianne Nussbaum, Isobel Tweedt, David A. Haake, Renee L. Galloway, Javier Monzón, Seth P.D. Riley, Jeff A. Sikich, Justin Brown, Anthony Friscia, Jessica W. Lynch, Katherine C. Prager, James O. Lloyd-Smith

**Author notes:** Disclaimer: The findings and conclusions in this report are those of the author(s) and do not necessarily represent the official position of the Centers for Disease Control and Prevention.

## Abstract

Leptospirosis is the most widespread zoonotic disease in the world, yet it is broadly understudied in multi-host wildlife systems. Knowledge gaps regarding *Leptospira* circulation in wildlife, particularly in densely populated areas, contribute to frequent misdiagnoses in humans and domestic animals. We assessed *Leptospira* prevalence levels and risk factors in five target wildlife species across the greater Los Angeles region: striped skunks (*Mephitis mephitis*), Northern raccoons (*Procyon lotor*), coyotes (*Canis latrans*), Virginia opossums (*Didelphis virginiana*), and fox squirrels (*Sciurus niger*). We sampled more than 960 individual animals, including over 700 from target species in the greater Los Angeles region, and an additional 260 sampled opportunistically from other regions and species. In the five target species, seroprevalences ranged from 5-60% and active infection prevalences ranged from 0.8-15.2% in all except fox squirrels (0%). Patterns of serologic reactivity suggest that mainland terrestrial wildlife, particularly mesocarnivores, could be the source of repeated observed introductions of *Leptospira* into local marine and island ecosystems. Overall, we found evidence of widespread *Leptospira* exposure in wildlife across Los Angeles and surrounding regions. This indicates exposure risk for humans and domestic animals and highlights that this pathogen can circulate endemically in many wildlife species even in densely populated urban areas.

## INTRODUCTION

Leptospirosis, the disease caused by pathogenic bacteria from the genus *Leptospira*, is the most widespread zoonotic disease in the world^1–3^. *Leptospira* is among the most generalist pathogens known and is capable of infecting a broad range of primarily mammalian hosts^1,4^. It is considered an emerging pathogen of concern^5^, with one million human cases and 60,000 deaths estimated globally each year^6,7^, and many more cases likely undiagnosed^3^. Given its extensive global impact, it is surprising that the ecology and epidemiology of this zoonotic pathogen remains understudied^8^. Though many pathogens, including all zoonoses by definition, are capable of infecting multiple hosts, studying complex generalist pathogen systems remains a challenge, particularly in wildlife for which extensive, multi-host surveillance is required but resources and sample access are often limited^9,10^. *Leptospira* poses additional challenges since host-pathogen epidemiology differs widely across host species and infecting strains, and multiple strains can cocirculate in communities. Consequently, extensive knowledge gaps remain regarding the importance of many hosts for onward transmission of *Leptospira*, and thus the risks they pose to other host species. This underscores the importance of broad surveillance efforts to inform veterinary and public health efforts, particularly in densely populated cities where the potential for human-wildlife contact is high and *Leptospira* transmission dynamics remain poorly understood.

Awareness of *Leptospira* genetic diversity has increased in recent years, with pathogenic *Leptospira* bacteria now categorized into 40 species^3,11,12^. The broad array of pathogenic *Leptospira* strains are classified into over 250 serovars based on similarities in cell surface antigens and serologic reactivity^1^, with related serovars traditionally grouped into serogroups, though neither serovar nor serogroup are reliable predictors of *Leptospira* species^13^. Transmission typically occurs after leptospires are shed in the urine of infected hosts, leading to environmental contamination that can indirectly infect susceptible hosts via mucous membranes or abraded skin^14–17^. Following this contact, leptospires colonize the kidneys and other organs, which can cause pathology and associated clinical signs ranging from mild flu-like symptoms to fulminant multi-organ system failure and death^15,18^. Disease severity varies across serovars and host species, with some broad host-serovar associations. Historically, the literature has categorized *Leptospira* epidemiology into: (i) maintenance hosts (and host-adapted serovars) that are likely to exhibit asymptomatic and/or chronic infections and maintain circulation of the pathogen, or (ii) accidental hosts (and non-host-adapted serovars) that exhibit symptomatic infections and do not maintain pathogen circulation^5^. These dichotomous classifications have been challenged in recent years, and there is clear evidence that clinical manifestations range widely irrespective of population-level circulation of the pathogen^19–21^. Evidence of infection is typically determined from urine or kidney samples using either polymerase chain reaction (PCR) to detect *Leptospira* DNA or culture isolates to detect live infectious leptospires. The most widely used diagnostic test for *Leptospira* spp. is the microscopic agglutination test (MAT) ^1,22^, which tests serum for anti-*Leptospira* antibodies to assess past exposure. Serum MAT panels typically include multiple serovars known or suspected to circulate in the geographic region of interest. Epidemiological patterns vary across climates and locations, with greater attention paid to the high incidence of human leptospirosis observed in wet tropical regions, and less concern historically placed on urban transmission, especially in more developed countries^5,7^. Despite increasing reports of *Leptospira* in urban areas, including dramatic outbreaks linked to flooding^23^, significant knowledge gaps remain regarding the prevalence and transmission risk across a variety of potential hosts in urban environments.

As urbanization increases worldwide, with over half of the global human population now residing in urban areas, it is important to identify the impacts of this anthropogenic change on wildlife and human health^24,25^. Urbanization can influence wildlife pathogen dynamics and zoonotic disease transmission risk through a variety of mechanisms, such as changing wildlife resource use, increasing exposure to toxicants, and altering community structure and contact rates^26^. At the same time, high densities of humans and domestic animals in urban settings provide increased opportunity for cross-species contacts and possible spillover of infection^27^. Homeless and marginalized individuals in crowded urban areas may be at increased risk for zoonotic disease transmission due to poor sanitation and intensified contact with a variety of urban wildlife species, even in high-income countries where urban homelessness is a growing public health crisis^28^. Climate change can further amplify urban transmission risk for some zoonoses, including *Leptospira* for which higher flood risk promotes environmental conditions favorable for pathogen survival and transmission^23^. Since changes to transmission risk are pathogen- and host-specific, investigations into pathogen dynamics and host diversity are critical for urban disease management and risk assessment. This is especially true for *Leptospira*, as most studies in cities have focused primarily on rodents in high-density urban centers^6,29–32^, with much less attention placed on other urban-adapted wildlife that may play a significant role in the ecology of this important zoonosis^5,33,34^. Improved knowledge of *Leptospira* epidemiology in a greater range of susceptible host species, and the diversity of *Leptospira* strains they carry, would provide valuable context for medical and veterinary clinicians assessing illness in humans and domestic animals.

In the United States, many *Leptospira* infections in humans, domestic dogs, horses, and livestock are associated with spillover from wildlife^35–37^. Increasing reports of leptospirosis have been observed in domestic dogs^38,39^ and humans in some locations (e.g. Hawaii and California^40–42^), further highlighting the need for expanded surveillance in wildlife. A recent study concluded that exposure to *Leptospira* is common in a variety of wildlife across the country, though more regional work is needed to understand the risk this poses to the health of humans, domestic pets and livestock^43^. In California, one historical wildlife survey reported *Leptospira* exposure or active infections in multiple species, including coyotes (*Canis latrans*), Northern raccoons (*Procyon lotor*) and striped skunks (*Mephitis mephitis*)^44^, while other surveys detected *Leptospira* in black bears (*Ursus americanus*)^45^ and feral pigs (*Sus scrofa*)^46^. A recent survey in northern California reported *Leptospira* in multiple mesocarnivore and rodent species^34^. Both skunks and raccoons were identified as potential reservoir hosts for *Leptospira interrogans* serovar Pomona^47^, a serovar which has also impacted California Channel Island terrestrial mammals and marine mammals along the California coast^20,21,48–52^.

In contrast with northern and coastal regions, there has been little surveillance for *Leptospira* in southern California. The diverse landscape of greater Los Angeles features a range of environments, from natural areas and agricultural land to a dense urban center, and many wildlife species that live alongside the ten million people in the region are potential *Leptospira* carriers^53^. There are a few reports of *Leptospira* exposure in some local animals, including dogs^54^, deer (*Odocoileus hemionus*)^55^, mountain lions (*Puma concolor*) and bobcats (*Lynx rufus*)^56^, but prior studies have had very limited host, geographic, and temporal ranges, and small sample sizes. The paucity of region-specific surveillance data in greater Los Angeles, the second largest metropolis in the United States, represents a critical public health gap, and more comprehensive wildlife screening is needed to assess zoonotic spillover risk from the full range of susceptible wildlife hosts.

We conducted the first in-depth surveillance of *Leptospira interrogans* in wildlife in the greater Los Angeles area to shed light on the prevalence and circulation of this multihost pathogen across a densely populated and complex urban landscape. We assessed *Leptospira* exposure and active infections in five common wildlife species: striped skunks, Northern raccoons, coyotes, Virginia opossums (*Didelphis virginiana*), and fox squirrels (*Sciurus niger*). We opportunistically sampled additional non-target species and regions to complement existing knowledge of this pathogen in California. We examined patterns in serologic reactivity across species to gain insights into multi-host *Leptospira* ecology and investigated how demographic and environmental conditions may impact *Leptospira* exposure across a heterogenous urbanization matrix. Our assessment of this zoonotic pathogen in the wildlife of greater Los Angeles has public health implications for millions of people and will inform veterinary and wildlife management agencies responsible for the health of humans and animals across the region.

## METHODS

### Study Animals

This study focused primarily on wildlife from the greater Los Angeles region in southern California. Opportunistic sample collection was approved by the California Department of Fish and Wildlife under scientific collecting permits SC-13267 & SC-13700 and took place from September 2015 to June 2020. Mountain lion samples were collected as part of the National Park Service mountain lion study, with capture and handling procedures permitted through the California Department of Fish and Wildlife (scientific collection permit number 5636) and the National Park Service Institutional Animal Care and Use Committee. For the purposes of this study, the greater Los Angeles region refers to Los Angeles County and surrounding counties: Orange, Riverside, San Bernardino, and Ventura. Sample collection focused on the following five common mammals in the Los Angeles region: striped skunk, Northern raccoon, coyote, Virginia opossum, and fox squirrel. These five species in greater Los Angeles will be referred to as our target species (Table 1, Fig. 1). Collaborating wildlife agencies donated carcasses or existing samples, with the majority coming from deceased animals following vehicle collisions, planned wildlife removal, or animals that were euthanized by animal control or rehabilitation agencies due to illness or injury. Carcasses were necropsied immediately or frozen at −20°C and thawed in a refrigerator prior to necropsy. Animal measurements and demographic information were collected at the time of necropsy, with age class (adult or juvenile) determined using a combination of animal size and tooth wear^**57**^.

**Table 1:**
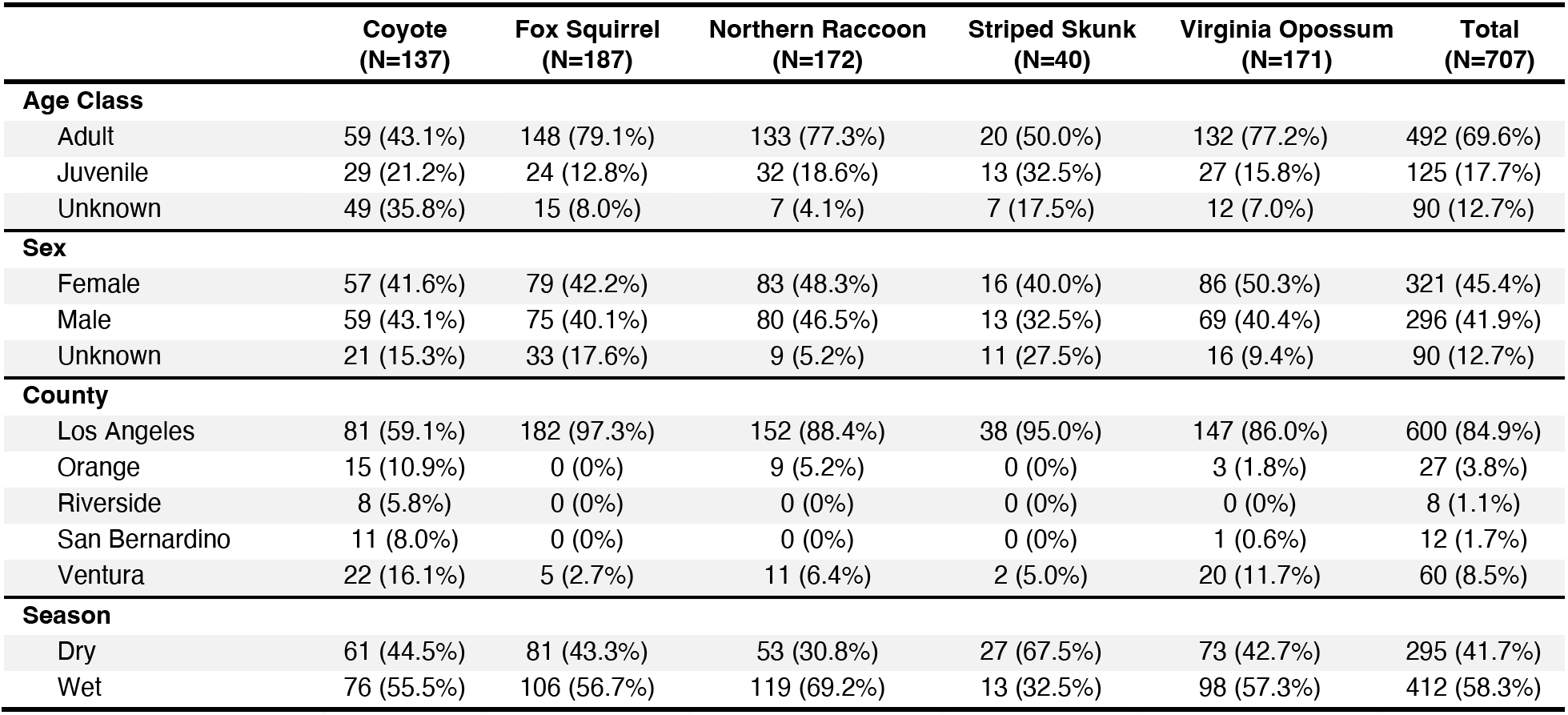
Descriptive characteristics for our five target wildlife species. Within-group percentages are proportions of column totals.

|  | Coyote<br>(N=137) | Fox Squirrel<br>(N=187) | Northern Raccoon<br>(N=172) | Striped Skunk<br>(N=40) | Virginia Opossum<br>(N=171) | Total<br>(N=707) |
| --- | --- | --- | --- | --- | --- | --- |
| **Age Class** |  |  |  |  |  |  |
| Adult | 59 (43.1%) | 148 (79.1%) | 133 (77.3%) | 20 (50.0%) | 132 (77.2%) | 492 (69.6%) |
| Juvenile | 29 (21.2%) | 24 (12.8%) | 32 (18.6%) | 13 (32.5%) | 27 (15.8%) | 125 (17.7%) |
| Unknown | 49 (35.8%) | 15 (8.0%) | 7 (4.1%) | 7 (17.5%) | 12 (7.0%) | 90 (12.7%) |
| **Sex** |  |  |  |  |  |  |
| Female | 57 (41.6%) | 79 (42.2%) | 83 (48.3%) | 16 (40.0%) | 86 (50.3%) | 321 (45.4%) |
| Male | 59 (43.1%) | 75 (40.1%) | 80 (46.5%) | 13 (32.5%) | 69 (40.4%) | 296 (41.9%) |
| Unknown | 21 (15.3%) | 33 (17.6%) | 9 (5.2%) | 11 (27.5%) | 16 (9.4%) | 90 (12.7%) |
| **County** |  |  |  |  |  |  |
| Los Angeles | 81 (59.1%) | 182 (97.3%) | 152 (88.4%) | 38 (95.0%) | 147 (86.0%) | 600 (84.9%) |
| Orange | 15 (10.9%) | 0 (0%) | 9 (5.2%) | 0 (0%) | 3 (1.8%) | 27 (3.8%) |
| Riverside | 8 (5.8%) | 0 (0%) | 0 (0%) | 0 (0%) | 0 (0%) | 8 (1.1%) |
| San Bernardino | 11 (8.0%) | 0 (0%) | 0 (0%) | 0 (0%) | 1 (0.6%) | 12 (1.7%) |
| Ventura | 22 (16.1%) | 5 (2.7%) | 11 (6.4%) | 2 (5.0%) | 20 (11.7%) | 60 (8.5%) |
| **Season** |  |  |  |  |  |  |
| Dry | 61 (44.5%) | 81 (43.3%) | 53 (30.8%) | 27 (67.5%) | 73 (42.7%) | 295 (41.7%) |
| Wet | 76 (55.5%) | 106 (56.7%) | 119 (69.2%) | 13 (32.5%) | 98 (57.3%) | 412 (58.3%) |

**Figure 1:**
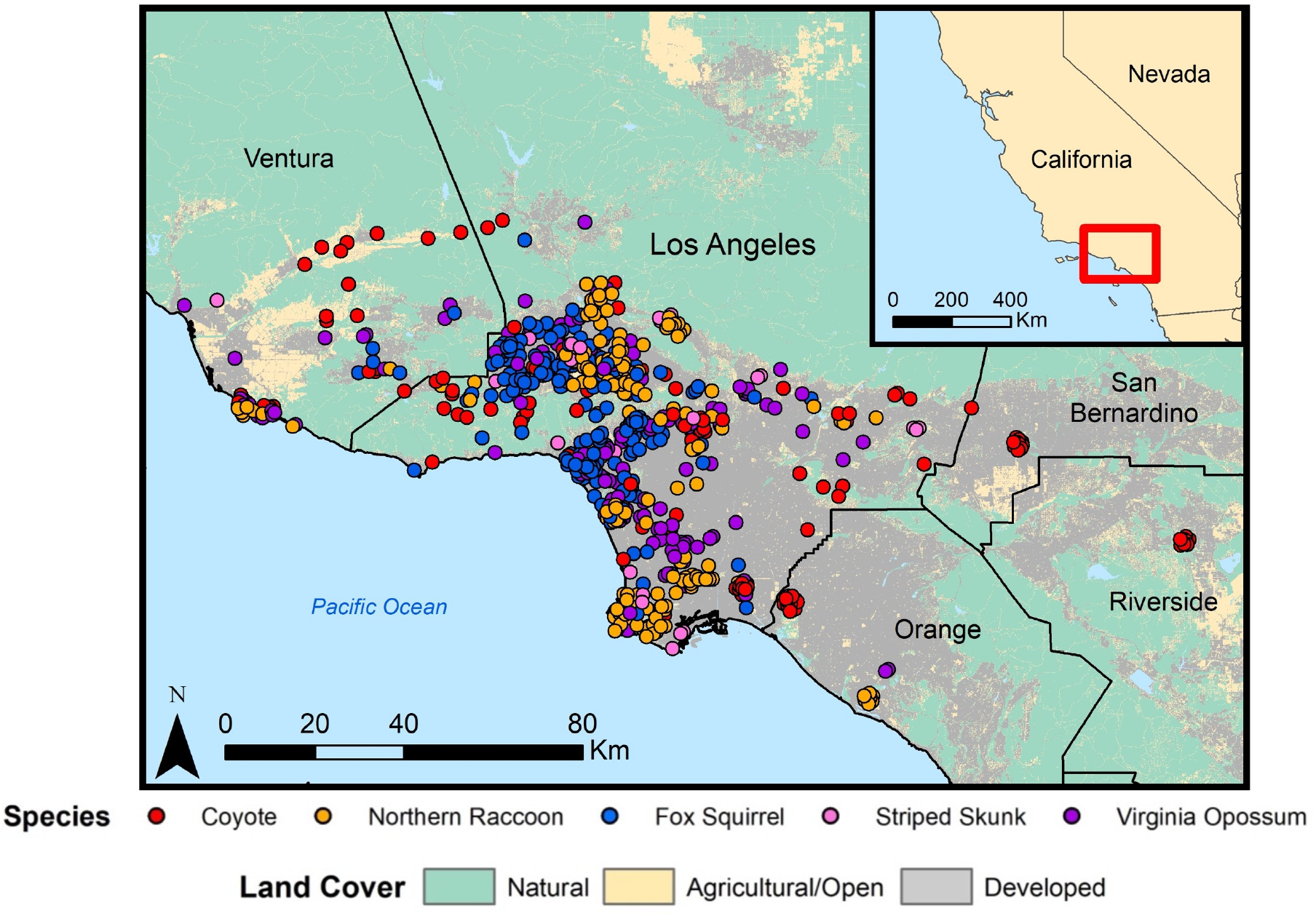
Distribution of *Leptospira* sample locations for the five target wildlife species in the greater Los Angeles region (2015-2020). Land cover data were obtained from the National Land Cover Database (2019).

### Sample Collection

Serum and urine samples from external agencies were analyzed and included when available. In fresh carcasses, intracardiac blood was collected into serum separation tubes, then kept in a cooler with an ice pack until centrifugation (1350 x g for 10-15 minutes). Kidney samples were collected from all animals that underwent a necropsy, and urine was collected when available using cystocentesis. The largest possible kidney sample that would fit in a 58 ml Whirl-Pak® was excised (approximate size: entire kidney from smaller mammals such as squirrels or half a kidney from larger mammals such as coyotes) and homogenized in the sealed Whirl-Pak® using manual pressure. Serum and urine samples were transferred into cryovials prior to storage, and all cryovials and Whirl-Paks® were stored at −20°C or −80°C prior to testing (−80°C preferred and transferred when possible).

### Leptospira Serology

Past exposure to *Leptospira* was assessed using microscopic agglutination testing (MAT). In this test, dark-field microscopy is used to assess the presence of anti-*Leptospira* antibodies in serum by evaluating agglutination (i.e. clumping) when samples are combined with live cultures of *Leptospira* species^22^. Serum samples are tested at doubling dilutions, and the reported endpoint titers represent the highest dilution that achieved a 50% agglutination using the reference strain being tested. Samples from 2015-2017 were analyzed at the California Animal Health and Food Safety Laboratory (CAHFS; Davis, California, USA) against a panel of 6 serovars that are common in the United States^43,58^: Bratislava, Canicola, Grippotyphosa, Hardjo, Icterohaemorrhagiae, and Pomona. Samples from 2017-2019 were analyzed at the Centers for Disease Control and Prevention (CDC; Atlanta, Georgia, USA) using an expanded panel of 20 serovars: Alexi, Australis, Autumnalis, Ballum, Bataviae, Borinca, Bratislava, Canicola, Celledoni, Cynopteri, Djasiman, Georgia, Grippotyphosa, Icterohaemorrhagiae, Javanica, Mankarso, Pomona, Pyrogenes, Tarassovi and Wolffi. To assess consistency between the two laboratories, we compared the subset of wildlife samples analyzed at both laboratories (n=469), demonstrating 98.3% agreement in seropositivity and minor quantitative differences between titers.

### Leptospira PCR Analysis

Pathogenic *Leptospira* DNA was assessed in urine (collected by cystocentesis) and homogenized kidney samples. For all species except mountain lions, samples collected from 2015-2017 were analyzed at the Hollings Marine Laboratory (Charleston, South Carolina, USA) using a quantitative polymerase chain reaction (qPCR) assay targeting the LipL32 gene as detailed in Wu et al. (2014)^59^. Samples collected after 2017 were analyzed at Colorado State University Veterinary Diagnostic Laboratory (Denver, Colorado, USA) using the VetMAX™ qPCR Master Mix kit and the primers specified by Wu et al. (2014)^59^. Minor modifications to the protocol resulted in slightly higher sensitivity, but results across laboratories were predominantly consistent^60^. Samples that had a cycle threshold value less than 37 were considered PCR positive. Samples from mountain lions were analyzed at the California Animal Health and Food Safety Laboratory (Davis, California, USA) using their standard protocol. Individuals that were PCR positive were assumed to have active infections and were considered likely to present a transmission risk, though caution should be taken with this interpretation since it is possible for PCR to detect non-viable *Leptospira* DNA^59^.

### Data Analysis

Prevalence was estimated for *Leptospira* exposure and infections. For *Leptospira* serology, individuals could be reactive to multiple serovars, so seroprevalence was calculated for each species in two ways: proportion positive against any serovar, and proportion positive within each specific serovar. All 95% binomial confidence intervals were estimated using package ‘PropCIs’ in *R* version 3.6.1^61^. Additional analyses were done in *R*, and maps were created using ArcGIS version 10.8.2^62^.

We used logistic regression to explore potential predictors of *Leptospira* exposure (as indicated by an antibody titer of 1:100 or higher to any serovar). The following covariates were considered: age class, sex, season (wet Nov-April vs. dry May-Oct)^34^, and composite land category (developed vs. agricultural/open vs. natural). Antibody titers have been identified as significant predictors of *Leptospira* shedding and PCR status in California sea lions^21,63^. We therefore explored this association in raccoons, the only species with a sufficient number of paired antibody-PCR samples available (n=81). Since these data exhibited complete separation (i.e. a clear distinction between the two outcomes), we applied a Firth’s bias-reduced logistic regression, a penalized maximum likelihood approach which is effective in the presence of data separation^64^, using the ‘logistf’ package in *R* to assess the association between antibody titers and PCR data.

Land cover analysis was conducted as detailed in Adducci II et al. (2020)^65^ using land cover data from the National Land Cover Database^66^. Sample localities occurred across a diverse landscape gradient, ranging from natural vegetation and open space to the urban core of Los Angeles (Fig. 1). To account for land cover variation within home ranges, we extracted 2019 land cover data (30m x 30m resolution) from home range buffers around each georeferenced sampling locality using the ‘raster’ and ‘rgdal’ packages in *R*. Buffer size varied by species based on home range estimates previously reported in the literature: 5 km^2^ for coyotes^65^, 2 km^2^ for raccoons and striped skunks^67^, 1 km^2^ for opossums^68^, and 0.5 km^2^ for fox squirrels^69^. As detailed in Adducci II et al. (2020)^65^, we grouped land cover classifications into three composite categories: urban/suburban (land with 20-100% impervious surface cover), agricultural/open (land used for pastures, crops and open development with <20% impervious surface cover), and natural (shrubland, forest, grassland and wetland). We then calculated the relative proportions of these three land categories for each individual home range buffer, and we used ternary plots to show *Leptospira* exposure (i.e. presence of antibodies) relative to land categories (Supplementary Fig. S1).

### Additional Species & Regions

On an opportunistic basis, we also tested samples provided by other agencies that included non-target regions of California and non-target species, including ground squirrels (*Otospermophilus beecheyi*), desert cottontails (*Sylvilagus audubonii*), feral pigs, bobcats, mountain lions, gray foxes (*Urocyon cinereoargenteus*), and red foxes (*Vulpes vulpes*). A total of 263 additional samples were collected from September 2015 thru May 2020. This included non-target species in Los Angeles County (n=74) and Ventura County (n=23), and all species from the following additional counties: Napa (n=14), Monterey (n=85), San Luis Obispo (n=61), Santa Barbara (n=6; Supplementary Table S1).

## RESULTS

### Leptospira Serology to Detect Exposure

We detected evidence of *Leptospira* exposure in all five target species sampled in the greater Los Angeles region (Fig. 2, Table 2). We first considered overall seroprevalence in each species, calculated as the proportion positive against any serovar. Fox squirrels had the highest overall seroprevalence at 60.6% (n=66/109, 95% CI: 50.7-69.8; Table 2), However, titer levels were low in this species, and most seropositive squirrels had maximum titers of 1:100 or 1:200 (Table 3). Other species had lower seroprevalence levels; seropositivity in raccoons was 32.6% (n=31/95, 95% CI: 23.4-43.0), followed by striped skunks at 28.6% (n=6/21; 95% CI: 11.3-52.2), coyotes at 25.5% (n=14/55, 95% CI: 14.7-39.0), and opossums at 5.2% (n=5/97; 95% CI: 1.7-11.6).

**Figure 2:**
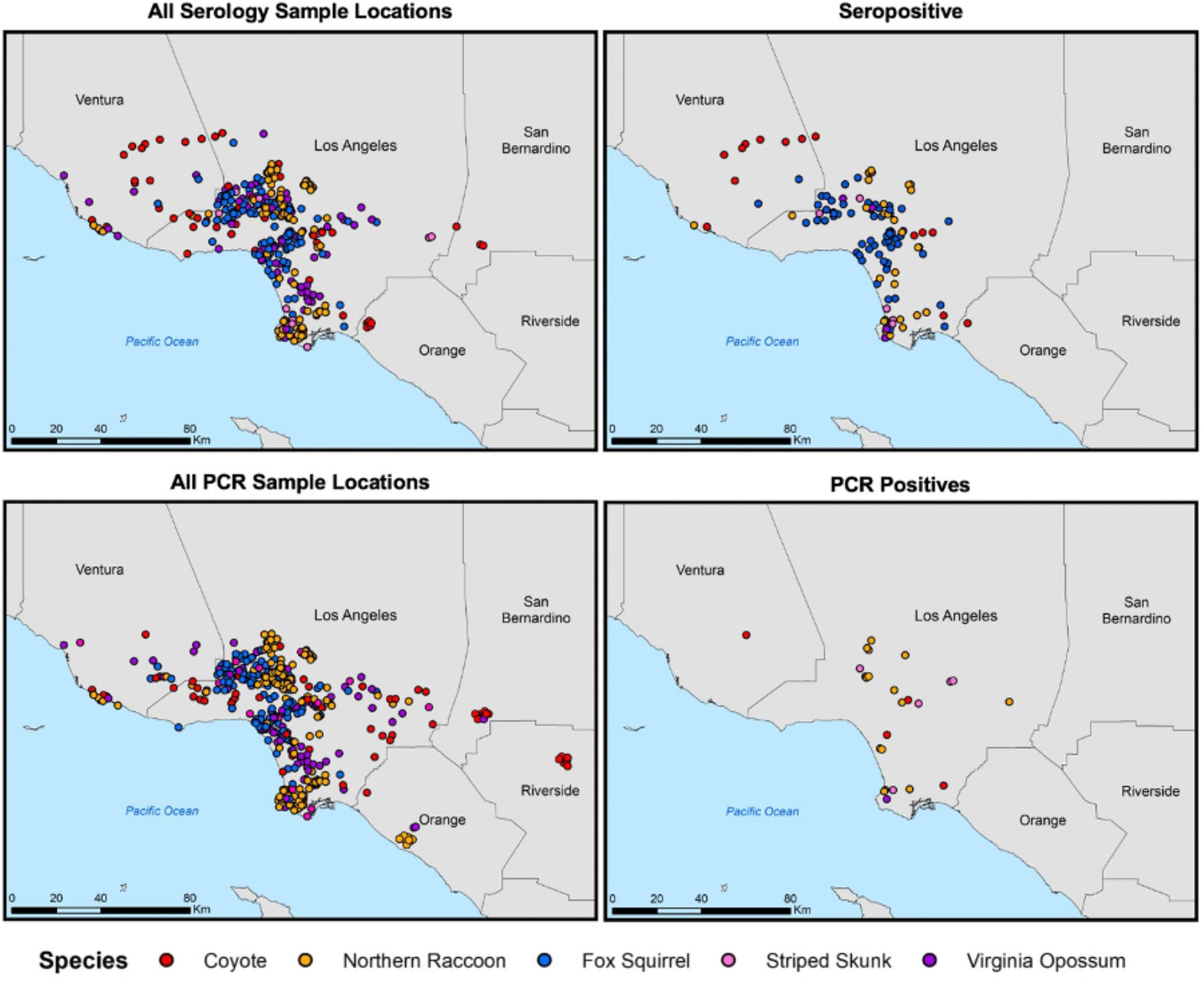
Locations of sample collection and associated *Leptospira* exposure and infection results for the five target wildlife species. Host sample locations are indicated in the left-hand panels, with the locations of animals with positive results shown in maps on the right. Samples tested by MAT for serum antibodies are shown in the top row, and those tested by PCR for *Leptospira* DNA are shown on the bottom row.

**Table 2:**
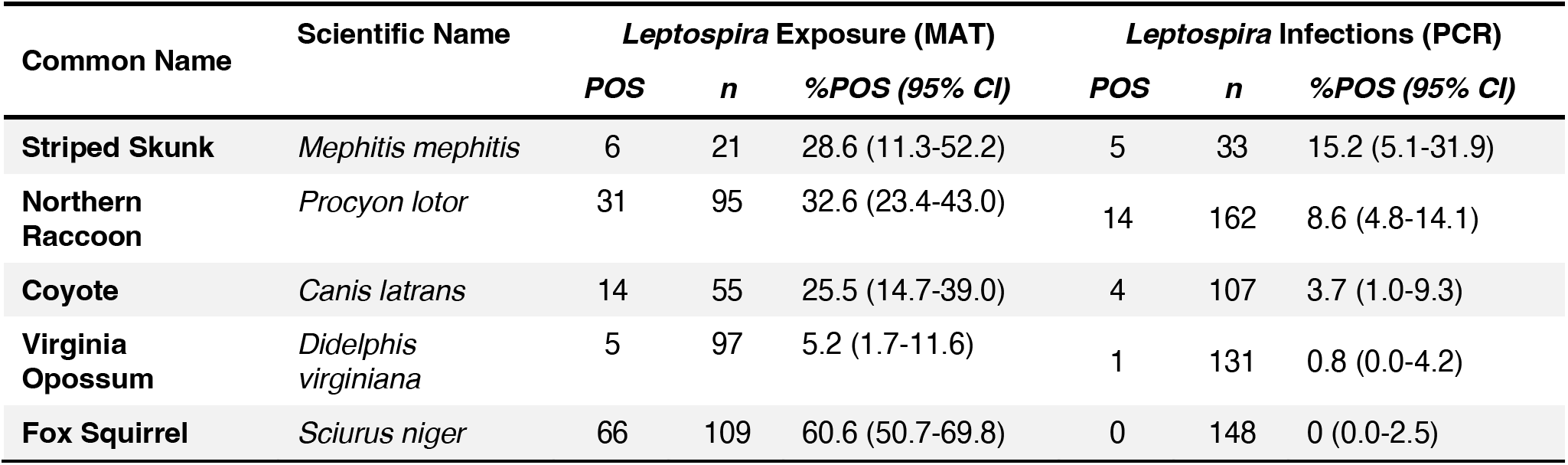
*Leptospira* exposure and infection by species. *Leptospira* antibody (MAT) and DNA (PCR) results in the five target species sampled in the greater Los Angeles region. Antibody results include seropositives to all serovars tested.

**Table 3:**
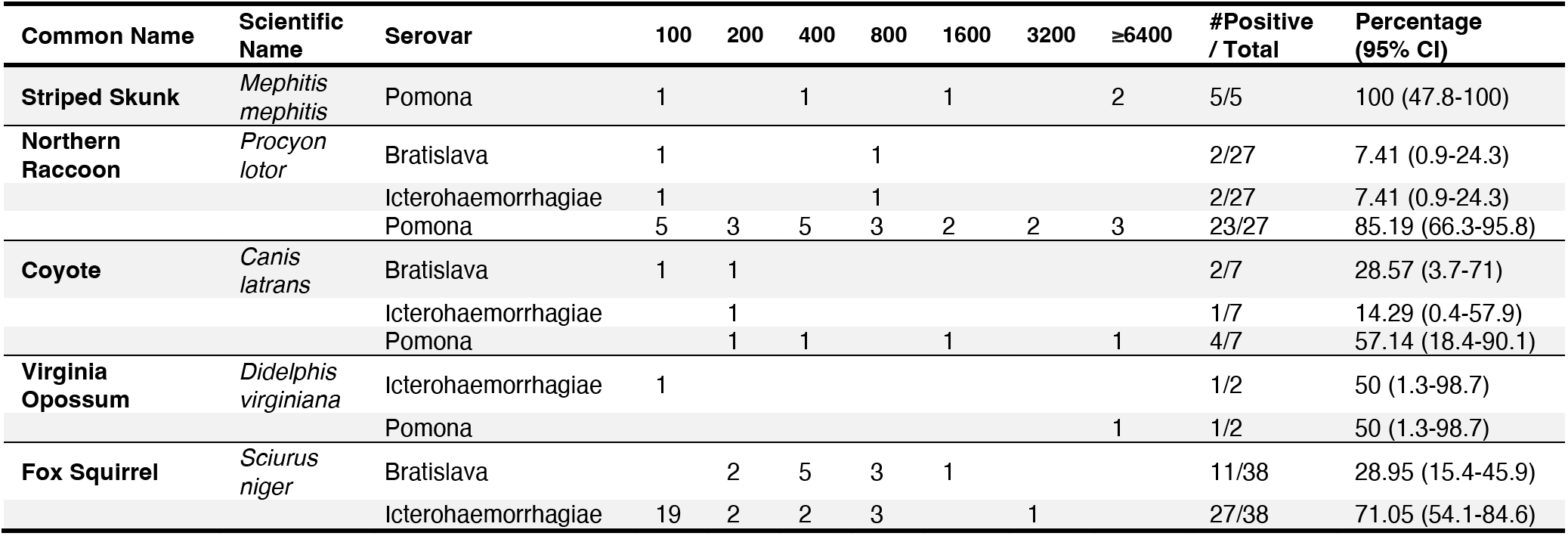
Maximum antibody titers for our five target wildlife species. Maximum antibody titers (MAT) and corresponding serovars for each individual from the five target species sampled in the greater Los Angeles region, reported for our five primary serovars (Bratislava, Canicola, Grippotyphosa, Icterohaemorrhagiae, and Pomona). Serovars that are not shown were never the maximum titer for that host species. In cases where there were ties for maximum titer, both serovars were counted in the table.

| Common Name | Scientific Name | Serovar | 100 | 200 | 400 | 800 | 1600 | 3200 | ≥6400 | #Positive / Total | Percentage (95% CI) |
| --- | --- | --- | --- | --- | --- | --- | --- | --- | --- | --- | --- |
| Striped Skunk | *Mephitis mephitis* | Pomona | 1 |  | 1 |  | 1 |  | 2 | 5/5 | 100 (47.8-100) |
| Northern Raccoon | *Procyon lotor* | Bratislava | 1 |  |  | 1 |  |  |  | 2/27 | 7.41 (0.9-24.3) |
|  |  | Icterohaemorrhagiae | 1 |  |  | 1 |  |  |  | 2/27 | 7.41 (0.9-24.3) |
|  |  | Pomona | 5 | 3 | 5 | 3 | 2 | 2 | 3 | 23/27 | 85.19 (66.3-95.8) |
| Coyote | *Canis latrans* | Bratislava | 1 | 1 |  |  |  |  |  | 2/7 | 28.57 (3.7-71) |
|  |  | Icterohaemorrhagiae |  | 1 |  |  |  |  |  | 1/7 | 14.29 (0.4-57.9) |
|  |  | Pomona |  | 1 | 1 |  | 1 |  | 1 | 4/7 | 57.14 (18.4-90.1) |
| Virginia Opossum | *Didelphis virginiana* | Icterohaemorrhagiae | 1 |  |  |  |  |  |  | 1/2 | 50 (1.3-98.7) |
|  |  | Pomona |  |  |  |  |  |  | 1 | 1/2 | 50 (1.3-98.7) |
| Fox Squirrel | *Sciurus niger* | Bratislava |  | 2 | 5 | 3 | 1 |  |  | 11/38 | 28.95 (15.4-45.9) |
|  |  | Icterohaemorrhagiae | 19 | 2 | 2 | 3 |  | 1 |  | 27/38 | 71.05 (54.1-84.6) |

To gain insight into the *Leptospira* variants generating these overall exposure levels, we first investigated patterns for the five main serovars that were tested for at both laboratories (serovars Bratislava, Canicola, Grippotyphosa, Icterohaemorrhagiae, and Pomona). We examined all positive MAT results to determine the full range of titer magnitude, the frequency of antibody detection against specific serovars, and the presence of antibody cross-reactivity within each of the five target host species (Fig. 3, Supplementary Table S2). All host species were serologically reactive to multiple serovars, and reactivity against serovar Pomona was detected in all species (Fig. 3). We then determined the serovar with the maximum MAT titer in each individual, as the best available indication of possible infecting serovar in the absence of isolates (Table 3; André-Fontaine & Triger, 2018). In skunks, raccoons, and coyotes, the highest antibody titers were most frequently against serovar Pomona (100%, 85%, and 57%, respectively; Table 3). Fox squirrels exhibited a pattern that was distinct from other hosts, with highest titers most often to serovar Icterohaemorrhagiae (71%), which was not highly reactive in any of the other host species. Of the two opossums that were reactive to this panel, one individual (the only opossum in this study with an active infection) had a maximum titer to serovar Pomona (1:12800). We emphasize that these maximum titer patterns do not give definitive information on the infecting serovar; confirmation requires serotyping of isolates.

**Figure 3:**
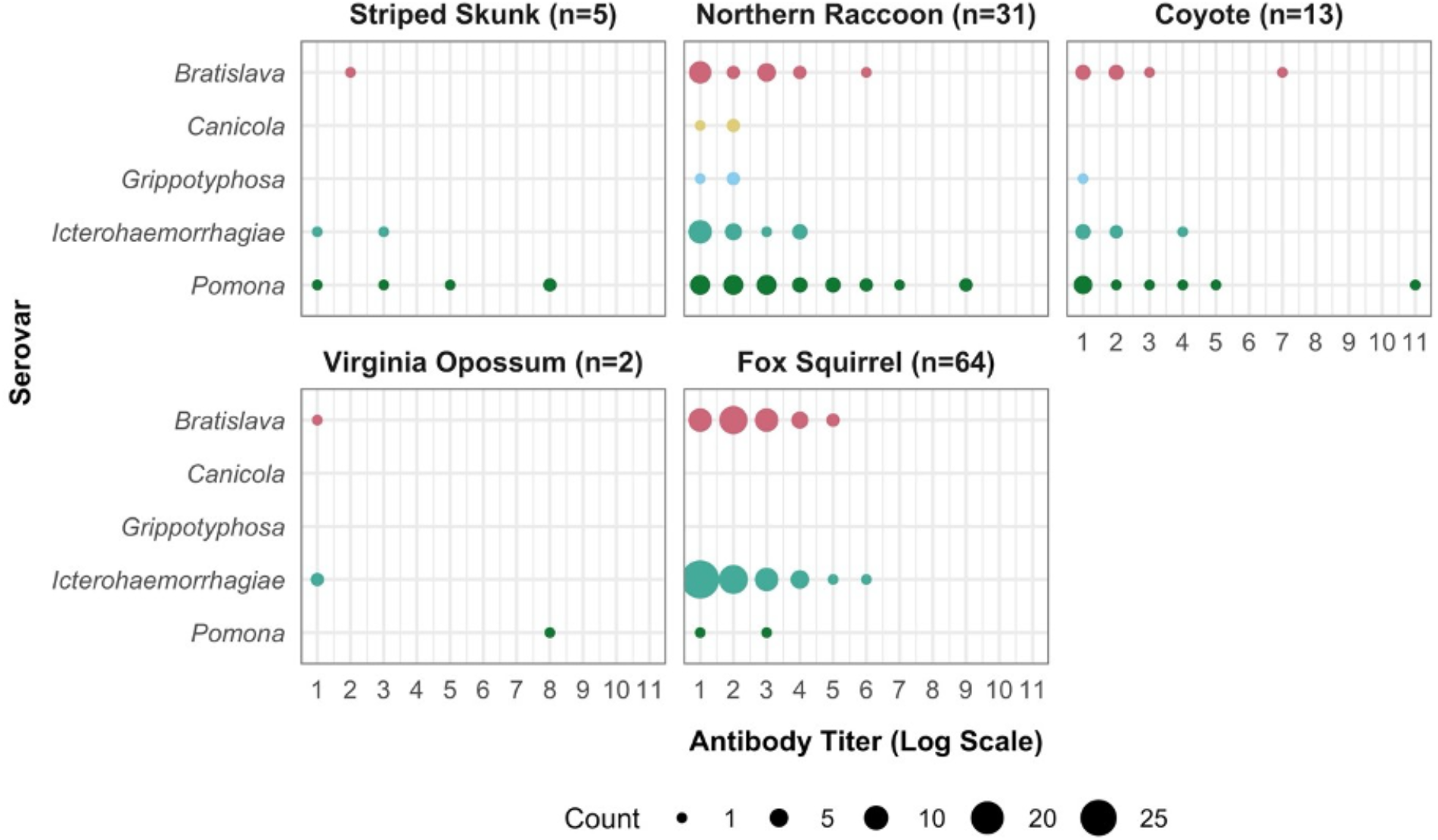
Samples positive for antibodies shown by host species, serovar, and titer level. Positive antibody results (MAT) for our five main serovars. Antibody titer (x-axis) is shown on a log scale (1:100 equivalent to 1, 1:200 equivalent to 2, 1:400 equivalent to 3, etc.).

We then considered MAT results for all 25 tested serovars across all species and regions (Supplementary Table S2). The serovar exhibiting highest seroprevalence varied by species, with skunks, raccoons, coyotes, opossums and fox squirrels testing positive most often to serovars Autumnalis (37.5%; n=3/8), Pomona (40.3%; n=54/134), Autumnalis (26.1%; n=12/46), Hardjo (9.3%; n=7/75) and Hardjo (66.7%; n=34/51), respectively. Though Autumnalis was the most frequently positive serovar in skunks and coyotes, Pomona was not significantly lower. Serovar Pomona comprised a larger sample size and total that tested positive in both species, with 26.9% (n=7/26) and 18.9% (n=14/74) of skunks and coyotes testing positive, respectively. Coyote titers against serovars Pomona and Autumnalis were strongly correlated (Spearman’s rho = 0.81), consistent with patterns previously observed in domestic dogs^70^. In aggregate, the data from coyotes and skunks are consistent with serovar Pomona being the primary infecting strain. Seven opossums were reactive to serovar Hardjo (Supplementary Table S2), making this the most commonly positive serovar overall in this species, though Pomona was the highest titer observed (1:12800), and the peak titer in the one opossum with an active infection detected. Aside from the low titer reactions to serovar Hardjo that more than doubled the overall seroprevalence of opossums, the majority of our conclusions in target species did not change with the consideration of the expanded serovar panel, with Icterohaemorrhagiae still a dominant serovar in squirrels and Pomona still a dominant serovar in the other species (Supplementary Table S2).

### Leptospira PCR to Detect Active Infections

Based on PCR results, we identified active infections (i.e. PCR positives) in all species except fox squirrels, with infection prevalence consistently lower than corresponding seroprevalence levels for each species (Table 2, Fig. 2). No active infections were detected in fox squirrels, despite seroprevalence being highest in this species. Infection prevalence ranged from 0.8% in opossums (n=1/131; 95% CI: 0.0-4.2) to 15% in skunks (n=5/33; 95% CI: 5.1-31.9), with coyotes and raccoons both intermediate at 3.7% (n=4/107; 95% CI: 1.0-9.3) and 8.6% (n=14/162; 95% CI: 4.8-14.1), respectively.

### Associations Between Serology and PCR Results

To analyze the association between maximum antibody titer against any serovar and active infection, we compared paired results for MAT titer and PCR, for which only raccoons had a sufficient sample size (n=81). Of those 81 animals, 29 were seropositive, and 86% of seropositive animals (n=25/29) had maximum titers to serovar Pomona. We found that individual maximum titers were significant predictors of PCR status in this species (Firth’s logistic regression; p-value = 2.09 × 10^−9^), which agrees with previous studies showing that *Leptospira* MAT titers are effective predictors of PCR status in California sea lions^21,63^. Individual raccoons with titers above 1:1600 were predicted to be highly likely to be PCR positive, and therefore likely to present a transmission risk because of their potential to be actively shedding *Leptospira* (Fig. 4).

**Figure 4:**
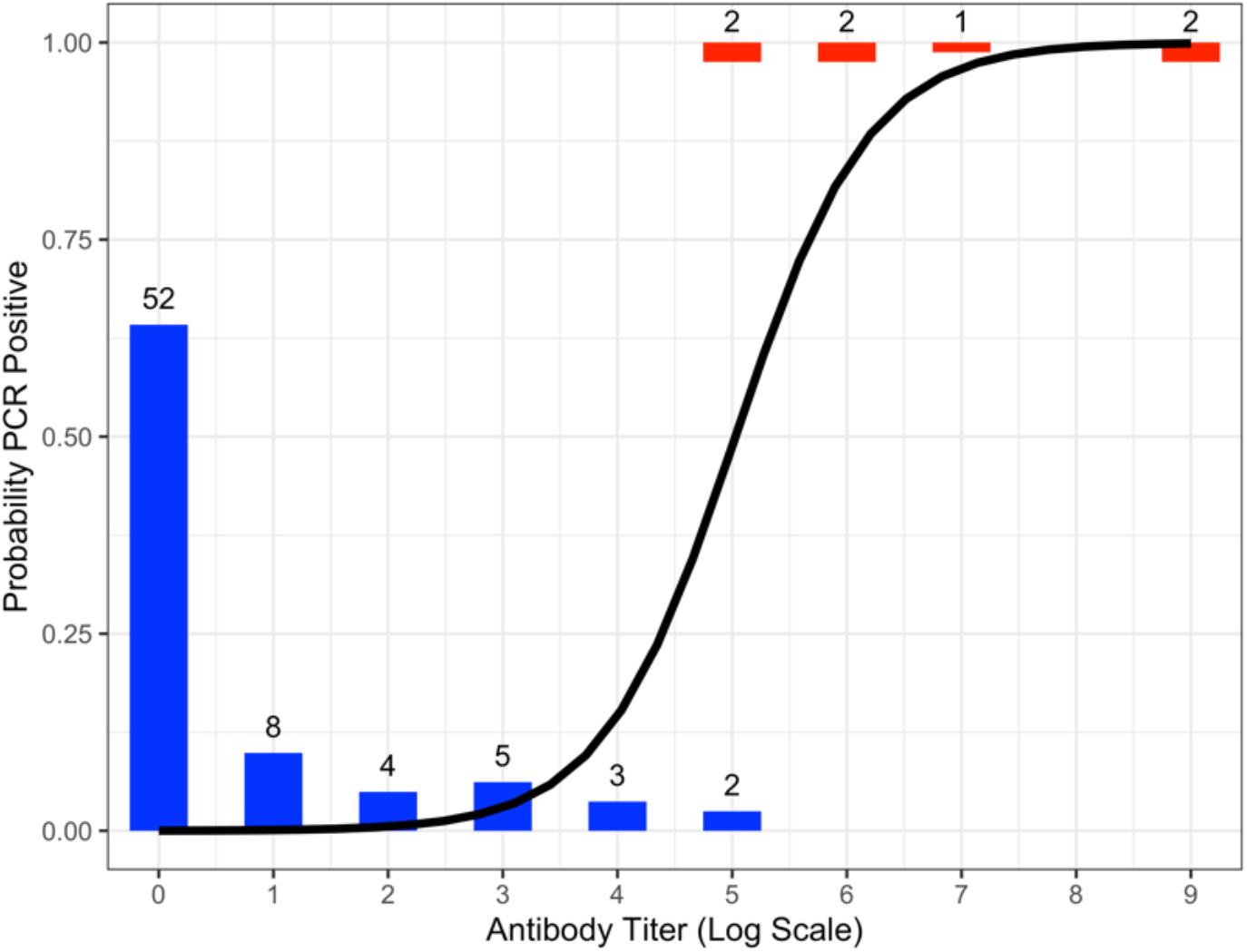
The predicted probability that raccoons are PCR-positive relative to maximum antibody titer to any serovar. The number of individuals with maximum MAT titer at a given level are represented by height of the bars and the corresponding number at the top of the bar. Animals with PCR negative results are shown in blue, whereas PCR positive individuals are shown in red. The solid black line shows the best-fitting solution of a Firth’s logistic regression, which predicts that individuals with MAT titers greater than 1:1600 (5 on the log scale) are at least 80% likely to be PCR positive.

### Spatial Patterns of Leptospira Exposure

We detected *Leptospira* exposure (i.e. MAT-positive individuals) throughout the sampled ranges of each host species (Fig. 2). When we evaluated sample location relative to the composite land cover classes, we found indications that different species use the landscape in different ways. For instance, fox squirrel and opossum samples were clustered around areas with higher levels of human development, providing evidence for increased use of urban and suburban regions in these non-native species (Supplementary Fig. S1). In contrast, coyotes and raccoons were found across all land classes. When we evaluated *Leptospira* exposure data in light of these land classes, no clear patterns emerged to distinguish the locations of positive and negative samples (Fig. 2, Supplementary Fig. S1), indicating that *Leptospira* circulates throughout the sampled range of each host species.

To probe how *Leptospira* exposure patterns are influenced by land cover, season and demographic factors, we used logistic regressions to assess possible correlates. This was done in all species except skunks, which were excluded due to small overall sample size (n=21). Of the covariates explored here (age class, sex, season, and composite land category), none exhibited significant correlations with *Leptospira* exposure in any of the target species (all p-values > 0.12 in univariate analyses). This lack of correlation aligns with the lack of clustering seen in the ternary plots (Fig. S1), supporting that *Leptospira* is distributed throughout the sampled range of these species.

### Additional Species & Regions

The non-target dataset (n=263 from 11 species) was comprised of samples from nontarget counties (n=166) and species (n=155). Only three of the 91 animals tested by PCR were positive (2 mountain lions and 1 feral pig), while antibodies were detected in 30.6% of non-target animals (n=63/206), with seroprevalence levels in individual species ranging from 0% to 60% (Supplementary Table S1). Feral pigs, sampled in Napa and San Luis Obispo (Supplementary Table S1), were more frequently reactive to serovars Bratislava (18.5%; n=10/54), Autumnalis (16.7%; n=9/54), Djasiman (14.8%; n=8/54) and Pomona (13.0%; n=7/54; Supplementary Table S2). Of the bobcats tested, 45% (n=5/11) were seropositive (Supplementary Table S1), with titers to Bratislava and Pomona most frequently positive (Supplementary Table S2). Of the non-target species in the greater Los Angeles region, 10% (n=3/30) of desert cottontails were seropositive, with the three positive individuals exhibiting titers against serovars Georgia, Icterohaemorrhagiae and Pomona, respectively (Supplementary Table S2). Two of the mountain lions from the greater Los Angeles region were PCR positive (18.2%; n=2/11), with both animals showing signs of clinical pathology (nephritis) upon necropsy.

## DISCUSSION

We conducted a large-scale survey of *Leptospira interrogans* in mainland terrestrial mammals in California, focusing on the understudied region of southern California. We had two goals: 1) to identify *Leptospira* prevalence, potential risk factors, and associated public health risks in the greater Los Angeles region, and 2) to assess serological patterns to inform our knowledge of broader multi-host circulation in coastal California wildlife. We identified *Leptospira* exposure in all target species sampled, and detected infections in all target species except fox squirrels. Widespread evidence of exposure, which was not concentrated around landscape or environmental factors, highlights that this pathogen is endemic and circulating throughout the ranges of these wildlife species in this major metropolitan area. Our overall findings extend the spatial and temporal scope of evidence that *Leptospira* is widespread in many wildlife species throughout California^34,44^.

In our five target species (skunks, raccoons, coyotes, opossums, and fox squirrels), we detected low to moderate levels of *Leptospira* infection and markedly higher seroprevalence levels. Skunks, raccoons and coyotes all exhibited moderate levels of infection, while infection prevalence was low in opossums and fox squirrels. Only one opossum was PCR positive, consistent with prior *Leptospira* surveys of opossums elsewhere in California, which have found no^71^ or low infection prevalence^34^. Fox squirrels had the highest seroprevalence of all our target hosts, though no infections were detected in this species. This could be due to a shorter duration of carriage, longer duration of titer decay (and hence seropositivity), or potentially an alternate route of transmission (e.g. sexual) and associated tissue distribution which could explain the lack of detection in the urinary tract. Seroprevalence levels in skunks, raccoons and coyotes were higher than opossums but lower than fox squirrels.

We then assessed seroreactivity patterns across all target and non-target species to better understand *Leptospira* ecology within our study system and possible connections to the broader California ecosystem. We focused on which serovar had the highest titer, though we emphasize that serology cannot conclusively identify the *Leptospira* strain causing an infection^72^. Coyotes, raccoons, and skunks typically had maximum antibody titers against serovar Pomona, consistent with findings in northern California where serovar Pomona predominated in these mesocarnivores^34^. Conversely, squirrels showed minimal seroreactivity to serovar Pomona and typically had low maximum titers against serovar Icterohaemorrhagiae, which was not highly reactive in other species. In addition to the widespread exposure detected in our target species, we detected exposure in all non-target species except gray foxes, and we confirmed that additional species (e.g. feral pigs and desert cottontails) can carry *Leptospira* in California.

Our study found lower prevalence of infection in our five target species than previously reported for the same species in more northern regions of California; comparing seroprevalence, only fox squirrels appeared to differ, with apparently higher seroprevalence in southern California^34^ (Supplementary Table S3). This may reflect true regional differences, though comparisons across studies should be considered carefully, owing to differences between laboratory assay protocols. For example, the prior study used a PCR cycle threshold of 45 to define positivity^34^, which results in higher sensitivity and lower specificity (i.e. a lower risk of false negatives, but a higher risk of false positives) than the cycle threshold of 37 used in this study. However, we expect quantitative differences to be relatively minor, and qualitative comparisons should still be valid across these studies. Regional differences in *Leptospira* infection incidence could also arise from environmental differences between northern and southern California. Landscape and environmental factors are known to impact *Leptospira*, as wet environmental conditions facilitate the bacteria’s survival and transmission. For example, higher rainfall has been associated with higher *Leptospira* incidence in domestic dogs^73^, a trend which has also been noted in the wetter region of northern California^74^. Though a seasonal *Leptospira* pattern was not observed in our Los Angeles data or wildlife in other California regions^34^, more extensive state-wide surveillance could detect such environmental patterns if they are present.

Our finding of widespread *Leptospira* circulation, with indications that serovar Pomona may be predominant in multiple host species, has potential implications for ongoing research in marine and terrestrial island mammals in California^20,48–52,60,75^. Phylogenetic analyses of *Leptospira* genomes isolated from California sea lions (*Zalophus californianus*), northern elephant seals (*Mirounga angustirostris*), Channel Island foxes (*Urocyon littoralis*) and spotted skunks (*Spilogale gracilis amphiala*) show evidence of repeated introductions of new strains of *L. interrogans* serovar Pomona into the broader coastal ecosystem^51^. The source of the introductions to the marine ecosystem is unknown but mainland terrestrial mammals are one possibility. Seroreactivity patterns in several species in our dataset echo patterns observed in the terrestrial mammals on the Channel Islands, indicating that mainland terrestrial mammals may play a role in the better-studied circulation of serovar Pomona in the broader California coastal ecosystem^20,51,76^.

Throughout this study, we relied on serological data to draw inferences about circulating strains of *Leptospira*. However, these observations should be interpreted with caution due to the potential for cross-reactivity between serovars^77^. Over half of the animals in this study were seroreactive to multiple serovars (65%; n=120/184), and definitively identifying the infecting serovar would require serotyping of isolates. The overall seroreactivity patterns detected here align with prior findings that high reactivity to serovar Pomona is common in California wildlife^51,52^, particularly in mesocarnivores^34^. However, to definitively assess the relatedness of strains across host species, including the potential hosts of serovar Pomona in our study (i.e. raccoons, skunks and coyotes), genetic sequence analyses of *Leptospira* will be required. This would yield key insights into multi-host pathogen dynamics, including possible transmission links between the potential hosts of serovar Pomona in our work and other host species in California marine and Channel Island ecosystems^51,78^. Conventionally, genetic sequencing requires culturing *Leptospira* from host samples, a major barrier due to the difficulty of culturing this slow-growing bacteria, though new culture-free approaches show promise.

Information on wildlife disease occurrence is crucial to assessing wildlife and human health risks, but collecting samples to support broad-scale wildlife surveillance can be very challenging. New approaches to analyzing existing data and more accessible samples could provide valuable tools to scale up surveillance. For example, analyses based on the relationship between serum antibody titers and shedding status, which has previously not been well characterized in wildlife^43^, may provide a proxy for determining infection prevalence from readily accessible serum samples. Here we showed that antibody titers are strongly predictive of shedding status in raccoons. Similar results were found in California sea lions^21,63^, indicating potential for a robust general pattern. This could be a useful screening tool for wildlife in situations where PCR results are not available or feasible to obtain, but more investigation is needed into the generality of this relationship given the known potential for *Leptospira* to exhibit strain- or host-specific patterns. This approach may not apply in some species, such as squirrels where antibody responses are variable^79,80^, or opossums where some individuals fail to mount an antibody response and seronegative shedding is possible^34,81^.

Baseline knowledge about the prevalence of zoonotic infections in urban wildlife has important utility for public health efforts. Diagnosing early cases of leptospirosis in humans and domestic animals can be challenging due to non-specific clinical signs. Raising clinical awareness about epidemiological risk factors (e.g. the prevalence in sympatric host species) is therefore critical for facilitating accurate diagnostics and disease risk assessments^3^. Broad-scale surveillance efforts also contribute to knowledge of circulating variants, which can be used to rapidly identify or exclude potential sources of transmission. For example, in 2021 there was a leptospirosis outbreak in domestic dogs in Los Angeles County, caused by *Leptospira interrogans* serovar Canicola^82^. No wildlife species in our study had predominant maximum titers to serovar Canicola, and few individuals showed any reactivity at all against this serovar. This corroborates the Los Angeles County Department of Public Health’s conclusion that this outbreak did not originate from local wildlife, highlighting the importance of longitudinal wildlife surveillance in determining (and ruling out) the source of outbreaks caused by multi-host pathogens.

This study has some limitations which suggest directions for additional surveillance efforts and future research. Since the surveillance efforts were finite and only five species could be sampled extensively, there could be unobserved host species contributing to *Leptospira* persistence and transmission in the greater Los Angeles region. Identifying these cryptic contributors, sometimes referred to as ‘epidemiological dark matter’, remains a frontier in disease ecology and emphasizes the need for ongoing research to understand multi-host pathogen dynamics. We were primarily dependent on collaborating agencies for salvaged and opportunistically collected samples for pathogen testing. This study design led to limited sample sizes, with a degree of spatial clustering around collaborator facilities (particularly for northern raccoons), which could have reduced our ability to detect spatial patterns. Samples may also have been biased towards more developed areas, as higher numbers of roads could have increased the numbers of traffic related deaths, and higher human density could also increase the likelihood of a sick or dead animal being reported. Additionally, the composite land use categories used here may have masked finer-scale spatial associations, and our use of circular home range buffers could overlook important behavior patterns in habitat use. Further investigations utilizing individual home ranges and land use categories or other metrics of urbanization (e.g. population density) could reveal additional spatial relationships undetected in our analyses.

This study provides the first in-depth look at *Leptospira* ecology in terrestrial wildlife across the greater Los Angeles area. Expanded knowledge of this pathogen in southern California, including comparisons of prevalence levels and serological patterns across host species, provides key insights into multi-host pathogen dynamics and the potential for cross-species transmission, including from wildlife to humans and their pets. Evidence of *Leptospira* circulation in Los Angeles wildlife has been lacking, contributing to the perception that the pathogen does not pose a major risk in the area. Our study found evidence consistent with endemic circulation of at least two strains of *Leptospira* among our five target species. High levels of exposure and wide geographic distribution indicate that this pathogen is ubiquitous across the region, with active infection rates substantial enough to warrant concern and to recommend that domestic dogs in Los Angeles be vaccinated against this disease. Better understanding of *Leptospira* ecology in California wildlife is critical to the management of this widely circulating pathogen. This highlights the need for systematic, broad-scale research efforts to monitor this pathogen in wildlife, domestic animals, and humans, particularly in densely populated urban settings like the greater Los Angeles region where the risk of zoonotic transmission may be high.

## Data availability

All data used for this paper will be archived on Github (https://github.com/SarahHelman/Lepto_in_SoCal_urban_wildlife).

## REFERENCES

1. Adler, B. & de la Peña Moctezuma, A. Leptospira and leptospirosis. Vet. Microbiol. 140, 287–296 (2010).

2. Fouts, D. E. et al. What Makes a Bacterial Species Pathogenic?:Comparative Genomic Analysis of the Genus Leptospira. PLoS Negl. Trop. Dis. 10, e0004403 (2016).

3. Sykes, J. E., Reagan, K. L., Nally, J. E., Galloway, R. L. & Haake, D. A. Role of Diagnostics in Epidemiology, Management, Surveillance, and Control of Leptospirosis. Pathogens 11, 395 (2022).

4. Cilia, G., Bertelloni, F., Albini, S. & Fratini, F. Insight into the Epidemiology of Leptospirosis: A Review of Leptospira Isolations from “Unconventional” Hosts. Animals 11, 191 (2021).

5. Levett, P. N. Leptospirosis. Clin. Microbiol. Rev. 14, 296–326 (2001).

6. Costa, F. et al. Patterns in Leptospira Shedding in Norway Rats (Rattus norvegicus) from Brazilian Slum Communities at High Risk of Disease Transmission. PLoS Negl. Trop. Dis. 9, e0003819 (2015).

7. Munoz-Zanzi, C. et al. A systematic literature review of leptospirosis outbreaks worldwide, 1970–2012. Rev. Panam. Salud Pública 44, 1 (2020).

8. Lloyd-Smith, J. O. et al. Epidemic Dynamics at the Human-Animal Interface. Science 326, 1362–1367 (2009).

9. Viana, M. et al. Assembling evidence for identifying reservoirs of infection. Trends Ecol. Evol. 29, 270–279 (2014).

10. Buhnerkempe, M. G. et al. Eight challenges in modelling disease ecology in multi-host, multi–agent systems. Epidemics 10, 26–30 (2015).

11. LPSN — List of Prokaryotic names with Standing in Nomenclature (bacterio.net). Genus Leptospira. https://lpsn.dsmz.de/genus/leptospira (2022).

12. Vincent, A. T. et al. Revisiting the taxonomy and evolution of pathogenicity of the genus Leptospira through the prism of genomics. PLoS Negl. Trop. Dis. 13, e0007270 (2019).

13. Levett, P. N. Systematics of Leptospiraceae. in Leptospira and Leptospirosis (ed. Adler, B.) vol. 387 11–20 (Springer Berlin Heidelberg, 2015).

14. Monahan, A. M., Callanan, J. J. & Nally, J. E. Proteomic Analysis of *Leptospira interrogans* Shed in Urine of Chronically Infected Hosts. Infect. Immun. 76, 4952–4958 (2008).

15. Haake, D. A. & Levett, P. N. Leptospirosis in Humans. in Leptospira and Leptospirosis (ed. Adler, B.) vol. 387 65–97 (Springer Berlin Heidelberg, 2015).

16. Casanovas-Massana, A. et al. Quantification of Leptospira interrogans Survival in Soil and Water Microcosms. Appl. Environ. Microbiol. 84, e00507–18 (2018).

17. Gostic, K. M. et al. Mechanistic dose-response modelling of animal challenge data shows that intact skin is a crucial barrier to leptospiral infection. Philos. Trans. R. Soc. Lond. B. Biol. Sci. 374, 20190367 (2019).

18. Ellis, W. A. Animal *Leptospirosis*. in Leptospira and Leptospirosis (ed. Adler, B.) 99–137 (Springer, 2015). doi: 10.1007/978-3-662-45059-8_6.

19. Ko, A. I., Goarant, C. & Picardeau, M. Leptospira: the dawn of the molecular genetics era for an emerging zoonotic pathogen. Nat. Rev. Microbiol. 7, 736–747 (2009).

20. Lloyd-Smith, J. O. et al. Cyclical changes in seroprevalence of leptospirosis in California sea lions: endemic and epidemic disease in one host species? BMC Infect. Dis. 7, 125–136 (2007).

21. Prager, K. C. et al. Linking longitudinal and cross-sectional biomarker data to understand host-pathogen dynamics: *Leptospira* in California sea lions (*Zalophus californianus*) as a case study. PLoS Negl. Trop. Dis. 14, e0008407 (2020).

22. Faine, S., Adler, B., Bolin, C. & Perolat, P. Leptospira and leptospirosis. (MediSci, 1999).

23. Lau, C. L., Smythe, L. D., Craig, S. B. & Weinstein, P. Climate change, flooding, urbanisation and leptospirosis: fuelling the fire? Trans. R. Soc. Trop. Med. Hyg. 104, 631–638 (2010).

24. Bradley, C. A. & Altizer, S. Urbanization and the ecology of wildlife diseases. Trends Ecol. Evol. 22, 95–102 (2007).

25. Hassell, J. M., Begon, M., Ward, M. J. & Fèvre, E. M. Urbanization and Disease Emergence: Dynamics at the Wildlife-Livestock-Human Interface. Trends Ecol. Evol. 32, 55–67 (2017).

26. Riley, S. P. D., Serieys, L. E. K. & Moriarty, J. G. Infectious Disease and Contaminants in Urban Wildlife: Unseen and Often Overlooked Threats. in Urban Wildlife (eds. McCleery, R. A., Moorman, C. E. & Peterson, M. N.) 175–215 (Springer US, 2014). doi: 10.1007/978-1-4899-7500-3_10.

27. Plowright, R. K. et al. Pathways to zoonotic spillover. Nat. Rev. Microbiol. 15, 502–510 (2017).

28. Leibler, J. H., Zakhour, C. M., Gadhoke, P. & Gaeta, J. M. Zoonotic and Vector-Borne Infections Among Urban Homeless and Marginalized People in the United States and Europe, 1990-2014. Vector Borne Zoonotic Dis. Larchmt. N 16, 435–444 (2016).

29. Vinetz, J. M., Glass, G. E., Flexner, C. E., Mueller, P. & Kaslow, D. C. Sporadic urban leptospirosis. Ann. Intern. Med. 125, 794–798 (1996).

30. Dupouey, J. et al. Human leptospirosis: an emerging risk in Europe? Comp. Immunol. Microbiol. Infect. Dis. 37, 77–83 (2014).

31. Minter, A. et al. A model for leptospire dynamics and control in the Norway rat (Rattus norvegicus) the reservoir host in urban slum environments. Epidemics 25, 26–34 (2018).

32. Boey, K., Shiokawa, K. & Rajeev, S. Leptospira infection in rats: A literature review of global prevalence and distribution. PLoS Negl. Trop. Dis. 13, e0007499 (2019).

33. Junge, R. E., Bauman, K., King, M. & Gompper, M. E. A serologic assessment of exposure to viral pathogens and Leptospira in an urban raccoon (Procyon lotor) population inhabiting a large zoological park. J. Zoo Wildl. Med. Off. Publ. Am. Assoc. Zoo Vet. 38, 18–26 (2007).

34. Straub, M. H. & Foley, J. E. Cross-sectional evaluation of multiple epidemiological cycles of *Leptospira* species in peri-urban wildlife in California. J. Am. Vet. Med. Assoc. 257, 840–848 (2020).

35. Davis, M. A. et al. Serological Survey for Antibodies to *Leptospira* in Dogs and Raccoons in Washington State. Zoonoses Public Health ???-??? (2008) doi: 10.1111/j.1863-2378.2008.01137.x.

36. Gautam, R., Wu, C.-C., Guptill, L. F., Potter, A. & Moore, G. E. Detection of antibodies against Leptospira serovars via microscopic agglutination tests in dogs in the United States, 2000–2007. J. Am. Vet. Med. Assoc. 237, 293–298 (2010).

37. Blessington, T., Schenck, A. P. & Levine, J. F. Frequency of Animal Leptospirosis in the Southern United States and the Implications for Human Health. South. Med. J. 113, 240–249 (2020).

38. White, A. M. et al. Hotspots of canine leptospirosis in the United States of America. Vet. J. 222, 29–35 (2017).

39. Smith, A. M., Stull, J. W. & Moore, G. E. Potential Drivers for the Re-Emergence of Canine Leptospirosis in the United States and Canada. Trop. Med. Infect. Dis. 7, 377 (2022).

40. Meites, E. et al. Reemerging Leptospirosis, California. Emerg. Infect. Dis. 10, 406–412 (2004).

41. Katz, A. R., Buchholz, A. E., Hinson, K., Park, S. Y. & Effler, P. V. Leptospirosis in Hawaii, USA, 1999-2008. Emerg. Infect. Dis. 17, 221–226 (2011).

42. Guerra, M. A. Leptospirosis: public health perspectives. Biol. J. Int. Assoc. Biol. Stand. 41, 295–297 (2013).

43. Pedersen, K. et al. Leptospira Antibodies Detected in Wildlife in the USA and the US Virgin Islands. J. Wildl. Dis. 54, 450–459 (2018).

44. Cirone, S. M., Riemann, H. P., Ruppanner, R., Behymer, D. E. & Franti, C. E. Evaluation of the hemagglutination test for epidemiologic studies of leptospiral antibodies in wild mammals. J. Wildl. Dis. 14, 193–202 (1978).

45. Ruppanner, R., Jessup, D. A., Ohishi, I., Behymer, D. E. & Franti, C. E. Serologic survey for certain zoonotic diseases in black bears in California. J. Am. Vet. Med. Assoc. 181, 1288–1291 (1982).

46. Clark, R. K., Jessup, D. A., Hird, D. W., Ruppanner, R. & Meyer, M. E. Serologic survey of California wild hogs for antibodies against selected zoonotic disease agents. J. Am. Vet. Med. Assoc. 183, 1248–1251 (1983).

47. Straub, M. H., Church, M., Glueckert, E. & Foley, J. E. Raccoons (*Procyon lotor*) and Striped Skunks (*Mephitis mephitis*) as Potential Reservoirs of *Leptospira* spp. in California. Vector-Borne Zoonotic Dis. 20, 418–426 (2020).

48. Gulland, F. M. D. et al. Leptospirosis in California sea lions (*Zalophus californianus*) stranded along the central California coast, 1981-1994. J. Wildl. Dis. 32, 572–580 (1996).

49. Greig, D. J., Gulland, F. M. D. & Kreuder, C. A Decade of Live California Sea Lion (*Zalophus californianus*) Strandings Along the Central California Coast: Causes and Trends, 1991-2000. Aquat. Mamm. 31, 11–22 (2005).

50. Colagross-Schouten, A. M., Mazet, J. A. K., Gulland, F. M. D., Miller, M. A. & Hietala, S. Diagnosis and seroprevalence of leptospirosis in California sea lions from coastal California. J. Wildl. Dis. 38, 7–17 (2002).

51. Mummah, R. O. *Leptospira* in the coastal California ecosystem: Challenges and solutions for analyzing complex wildlife disease data. (University of California Los Angeles, 2021).

52. Zuerner, R. L. et al. Geographical dissemination of Leptospira interrogans serovar Pomona during seasonal migration of California sea lions. Vet. Microbiol. 137, 105–110 (2009).

53. U.S. Census Bureau QuickFacts: Los Angeles city, California. https://www.census.gov/quickfacts/fact/table/losangelescountycalifornia,losangelescitycalifornia/PST045222,PST045221.

54. Greene, M. R. A survey for leptospirosis in Southern California. Am. J. Epidemiol. 34-SectionB, 87–90 (1941).

55. Roug, A., Swift, P., Torres, S., Jones, K. & Johnson, C. K. Serosurveillance for Livestock Pathogens in Free-Ranging Mule Deer (Odocoileus hemionus). PLoS ONE 7, e50600 (2012).

56. Straub, M. H., Rudd, J. L., Woods, L. W., Clifford, D. L. & Foley, J. E. Leptospira prevalence and its association with renal pathology in mountain lions (Puma concolor) and bobcats (lynx rufus) in California, USA. J. Wildl. Dis. 57, (2021).

57. Grau, G. A., Sanderson, G. C. & Rogers, J. P. Age Determination of Raccoons. J. Wildl. Manag. 34, 364 (1970).

58. Pedersen, K. et al. Widespread detection of antibodies to *Leptospira* in feral swine in the United States. Epidemiol. Infect. 143, 2131–2136 (2015).

59. Wu, Q. et al. Development of a real-time PCR for the detection of pathogenic *Leptospira* spp. in California sea lions. Dis. Aquat. Organ. 110, 165–172 (2014).

60. Lloyd-Smith, J. O. & Prager, K. C. Leptospirosis in endangered island foxes and California sea lions: Outbreak prediction and prevention in a changing world. https://apps.dtic.mil/sti/citations/AD1189863 (2021).

61. R Core Team. R: A language and environment for statistical computing. (2021).

62. ArcGIS Desktop. (2011).

63. Helman, S. K. From California sea lions to urban coyotes: Maximizing insights from Leptospira surveillance in coastal California wildlife. (UCLA, 2022).

64. Firth, D. Bias reduction of maximum likelihood estimates. Biometrika 80, 27–38 (1993).

65. Adducci II, A. et al. Urban coyotes are genetically distinct from coyotes in natural habitats. J. Urban Ecol. 6, juaa010 (2020).

66. US. Geological Survey (USGS). NLCD 2019 Land Cover Conterminous United States. (2019).

67. Šálek, M., Drahníková, L. & Tkadlec, E. Changes in home range sizes and population densities of carnivore species along the natural to urban habitat gradient. Mammal Rev. 45, 1–14 (2015).

68. Wright, J. D., Burt, M. S. & Jackson, V. L. Influences of an Urban Environment on Home Range and Body Mass of Virginia Opossums (*Didelphis virginiana*). Northeast. Nat. 19, 77–86 (2012).

69. Prince, A., DePerno, C. S., Gardner, B. & Moorman, C. E. Survival and Home-Range Size of Southeastern Fox Squirrels in North Carolina. Southeast. Nat. 13, 456 (2014).

70. Moore, G. E. et al. Canine Leptospirosis, United States, 2002–2004. Emerg. Infect. Dis. 12, 501–503 (2006).

71. Krueger, L. et al. Identification of Zoonotic and Vector-borne Infectious Agents Associated with Opossums (Didelphis virginiana) in Residential Neighborhoods of Orange County, California. Proc. Vertebr. Pest Conf. 27, (2016).

72. André-Fontaine, G. & Triger, L. MAT cross-reactions or vaccine cross-protection: retrospective study of 863 leptospirosis canine cases. Heliyon 4, e00869 (2018).

73. Ward, M. P. Seasonality of canine leptospirosis in the United States and Canada and its association with rainfall. Prev. Vet. Med. 56, 203–213 (2002).

74. Adin, C. A. & Cowgill, L. D. Treatment and outcome of dogs with leptospirosis: 36 cases (1990-1998). J. Am. Vet. Med. Assoc. 216, 371–375 (2000).

75. Buhnerkempe, M. G. et al. Detecting signals of chronic shedding to explain pathogen persistence: *Leptospira interrogans* in California sea lions. J. Anim. Ecol. 86, 460–472 (2017).

76. Prager, K. C. et al. Asymptomatic and chronic carriage of Leptospira interrogans serovar Pomona in California sea lions (Zalophus californianus). Vet. Microbiol. 164, 177–183 (2013).

77. Chirathaworn, C., Inwattana, R., Poovorawan, Y. & Suwancharoen, D. Interpretation of microscopic agglutination test for leptospirosis diagnosis and seroprevalence. Asian Pac. J. Trop. Biomed. 4, S162–164 (2014).

78. Borremans, B., Faust, C., Manlove, K. R., Sokolow, S. H. & Lloyd-Smith, J. O. Crossspecies pathogen spillover across ecosystem boundaries: mechanisms and theory. Philos. Trans. R. Soc. B Biol. Sci. 374, 20180344 (2019).

79. Diesch, S. L., Crawford, R. P., McCulloch, W. F. & Top, F. H. Human Leptospirosis Acquired from Squirrels. N. Engl. J. Med. 276, 838–842 (1967).

80. Dirsmith, K. et al. Leptospirosis in Fox Squirrels (Sciurus niger) of Larimer County, Colorado, USA. J. Wildl. Dis. 49, 641–645 (2013).

81. Reilly, J. R. The susceptibility of five species of wild animals to experimental infection with Leptospira grippotyphosa. J. Wildl. Dis. 6, 289–294 (1970).

82. LA County Department of Public Health (LADPH). Leptospirosis in Dogs in Los Angeles County in 2021. http://www.publichealth.lacounty.gov/vet/Leptospirosis2021.htm (2022).

