## Supplementary Tables and Figures for "Pathogenic *Leptospira* are widespread in the urban wildlife of southern California"

### Table S1: *Leptospira* test results from non-target species and regions.

| | Boar (N=54) | Bobcat (N=11) | Coyote (N=20) | Desert Cottontail (N=38) | Gray Fox (N=9) | Ground Squirrel (N=38) | Mountain Lion (N=11) | Northern Raccoon (N=40) | Red Fox (N=5) | Striped Skunk (N=5) | Virginia Opossum (N=32) | Total (N=263) |
|---|---|---|---|---|---|---|---|---|---|---|---|---|
| **MAT** | | | | | | | | | | | | |
| Negative | 39 (72.2%) | 6 (54.5%) | 12 (60.0%) | 27 (71.1%) | 4 (44.4%) | 2 (5.3%) | 1 (9.1%) | 16 (40.0%) | 3 (60.0%) | 3 (60.0%) | 30 (93.8%) | 143 (54.4%) |
| Positive | 15 (27.8%) | 5 (45.5%) | 7 (35.0%) | 3 (7.9%) | 0 (0%) | 2 (5.3%) | 1 (9.1%) | 24 (60.0%) | 2 (40.0%) | 2 (40.0%) | 2 (6.3%) | 63 (24.0%) |
| Not Tested | 0 (0%) | 0 (0%) | 1 (5.0%) | 8 (21.1%) | 5 (55.6%) | 34 (89.5%) | 9 (81.8%) | 0 (0%) | 0 (0%) | 0 (0%) | 0 (0%) | 57 (21.7%) |
| **PCR** | | | | | | | | | | | | |
| Negative | 7 (13.0%) | 1 (9.1%) | 2 (10.0%) | 27 (71.1%) | 6 (66.7%) | 36 (94.7%) | 9 (81.8%) | 0 (0%) | 0 (0%) | 0 (0%) | 0 (0%) | 88 (33.5%) |
| Positive | 1 (1.9%) | 0 (0%) | 0 (0%) | 0 (0%) | 0 (0%) | 0 (0%) | 2 (18.2%) | 0 (0%) | 0 (0%) | 0 (0%) | 0 (0%) | 3 (1.1%) |
| Not Tested | 46 (85.2%) | 10 (90.9%) | 18 (90.0%) | 11 (28.9%) | 3 (33.3%) | 2 (5.3%) | 0 (0%) | 40 (100%) | 5 (100%) | 5 (100%) | 32 (100%) | 172 (65.4%) |
| **Sex** | | | | | | | | | | | | |
| Female | 27 (50.0%) | 1 (9.1%) | 5 (25.0%) | 16 (42.1%) | 1 (11.1%) | 13 (34.2%) | 6 (54.5%) | 1 (2.5%) | 2 (40.0%) | 0 (0%) | 5 (15.6%) | 77 (29.3%) |
| Male | 25 (46.3%) | 2 (18.2%) | 8 (40.0%) | 10 (26.3%) | 5 (55.6%) | 23 (60.5%) | 5 (45.5%) | 2 (5.0%) | 2 (40.0%) | 3 (60.0%) | 3 (9.4%) | 88 (33.5%) |
| Unknown | 2 (3.7%) | 8 (72.7%) | 7 (35.0%) | 12 (31.6%) | 3 (33.3%) | 2 (5.3%) | 0 (0%) | 37 (92.5%) | 1 (20.0%) | 2 (40.0%) | 24 (75.0%) | 98 (37.3%) |
| **Age Class** | | | | | | | | | | | | |
| Adult | 35 (64.8%) | 7 (63.6%) | 14 (70.0%) | 36 (94.7%) | 6 (66.7%) | 25 (65.8%) | 10 (90.9%) | 38 (95.0%) | 4 (80.0%) | 5 (100%) | 31 (96.9%) | 211 (80.2%) |
| Juvenile | 19 (35.2%) | 2 (18.2%) | 4 (20.0%) | 0 (0%) | 2 (22.2%) | 9 (23.7%) | 1 (9.1%) | 2 (5.0%) | 1 (20.0%) | 0 (0%) | 0 (0%) | 40 (15.2%) |
| Unknown | 0 (0%) | 2 (18.2%) | 2 (10.0%) | 2 (5.3%) | 1 (11.1%) | 4 (10.5%) | 0 (0%) | 0 (0%) | 0 (0%) | 0 (0%) | 1 (3.1%) | 12 (4.6%) |
| **County** | | | | | | | | | | | | |
| Napa | 14 (25.9%) | 0 (0%) | 0 (0%) | 0 (0%) | 0 (0%) | 0 (0%) | 0 (0%) | 0 (0%) | 0 (0%) | 0 (0%) | 0 (0%) | 14 (5.3%) |
| San Luis Obispo | 40 (74.1%) | 0 (0%) | 9 (45.0%) | 0 (0%) | 0 (0%) | 0 (0%) | 0 (0%) | 1 (2.5%) | 4 (80.0%) | 3 (60.0%) | 4 (12.5%) | 61 (23.2%) |
| Los Angeles | 0 (0%) | 2 (18.2%) | 0 (0%) | 24 (63.2%) | 5 (55.6%) | 36 (94.7%) | 7 (63.6%) | 0 (0%) | 0 (0%) | 0 (0%) | 0 (0%) | 74 (28.1%) |
| Monterey | 0 (0%) | 5 (45.5%) | 7 (35.0%) | 0 (0%) | 2 (22.2%) | 1 (2.6%) | 0 (0%) | 39 (97.5%) | 1 (20.0%) | 2 (40.0%) | 28 (87.5%) | 85 (32.3%) |
| Santa Barbara | 0 (0%) | 1 (9.1%) | 4 (20.0%) | 0 (0%) | 1 (11.1%) | 0 (0%) | 0 (0%) | 0 (0%) | 0 (0%) | 0 (0%) | 0 (0%) | 6 (2.3%) |
| Ventura | 0 (0%) | 3 (27.3%) | 0 (0%) | 14 (36.8%) | 1 (11.1%) | 1 (2.6%) | 4 (36.4%) | 0 (0%) | 0 (0%) | 0 (0%) | 0 (0%) | 23 (8.7%) |
| **Season** | | | | | | | | | | | | |
| Dry | 17 (31.5%) | 3 (27.3%) | 8 (40.0%) | 16 (42.1%) | 3 (33.3%) | 28 (73.7%) | 8 (72.7%) | 13 (32.5%) | 2 (40.0%) | 0 (0%) | 6 (18.8%) | 104 (39.5%) |
| Wet | 28 (51.9%) | 8 (72.7%) | 11 (55.0%) | 22 (57.9%) | 6 (66.7%) | 10 (26.3%) | 3 (27.3%) | 27 (67.5%) | 3 (60.0%) | 5 (100%) | 26 (81.3%) | 149 (56.7%) |
| Unknown | 9 (16.7%) | 0 (0%) | 1 (5.0%) | 0 (0%) | 0 (0%) | 0 (0%) | 0 (0%) | 0 (0%) | 0 (0%) | 0 (0%) | 0 (0%) | 10 (3.8%) |

**Table S2: *Leptospira* serovars and antibody titer levels for all California wildlife samples.** A description of the total number tested by species and serovar (n), the number of animals that were positive to each titer level, and the row-wise positivity (% positive) with binomial confidence intervals (95% CI) are shown in the table below.

| Common Name | *Scientific Name* | n | Serovar | 100 | 200 | 400 | 800 | 1600 | 3200 | >=6400 | N | Prevalence (%) |
|---|---|---|---|---|---|---|---|---|---|---|---|---|
| **Feral Pigs** | *Sus scrofa* | 54 | Autumnalis | 4 | 1 | 3 | 0 | 0 | 0 | 0 | 9 | 16.67 (7.9-29.3) |
| | | 54 | Bratislava | 4 | 4 | 2 | 0 | 0 | 0 | 0 | 10 | 18.52 (9.3-31.4) |
| | | 54 | Canicola | 1 | 0 | 0 | 0 | 0 | 0 | 0 | 1 | 1.85 (0-9.9) |
| | | 54 | Cynopteri | 1 | 1 | 0 | 0 | 0 | 0 | 0 | 2 | 3.7 (0.5-12.7) |
| | | 54 | Djasiman | 3 | 1 | 3 | 1 | 0 | 0 | 0 | 8 | 14.81 (6.6-27.1) |
| | | 54 | Georgia | 0 | 1 | 0 | 0 | 0 | 0 | 0 | 1 | 1.85 (0-9.9) |
| | | 54 | Icterohaemorrhagiae | 2 | 0 | 1 | 0 | 0 | 0 | 0 | 3 | 5.56 (1.2-15.4) |
| | | 54 | Mankarso | 1 | 0 | 1 | 0 | 0 | 0 | 0 | 2 | 3.7 (0.5-12.7) |
| | | 54 | Pomona | 2 | 0 | 1 | 0 | 1 | 1 | 2 | 7 | 12.96 (5.4-24.9) |
| | | 54 | Pyrogenes | 0 | 1 | 0 | 0 | 0 | 0 | 0 | 1 | 1.85 (0-9.9) |
| | | 54 | Tarassovi | 1 | 0 | 0 | 0 | 0 | 0 | 0 | 1 | 1.85 (0-9.9) |
| **Bobcat** | *Lynx rufus* | 7 | Autumnalis | 0 | 0 | 0 | 0 | 0 | 1 | 0 | 1 | 14.29 (0.4-57.9) |
| | | 7 | Bataviae | 1 | 0 | 0 | 0 | 0 | 0 | 0 | 1 | 14.29 (0.4-57.9) |
| | | 11 | Bratislava | 2 | 1 | 0 | 1 | 0 | 0 | 0 | 4 | 36.36 (10.9-69.2) |
| | | 11 | Canicola | 0 | 0 | 0 | 0 | 1 | 0 | 0 | 1 | 9.09 (0.2-41.3) |
| | | 7 | Cynopteri | 1 | 0 | 0 | 0 | 0 | 0 | 0 | 1 | 14.29 (0.4-57.9) |
| | | 7 | Djasiman | 0 | 0 | 0 | 0 | 0 | 1 | 0 | 1 | 14.29 (0.4-57.9) |
| | | 11 | Grippotyphosa | 0 | 0 | 0 | 0 | 1 | 0 | 0 | 1 | 9.09 (0.2-41.3) |
| | | 11 | Icterohaemorrhagiae | 0 | 1 | 0 | 0 | 0 | 1 | 0 | 2 | 18.18 (2.3-51.8) |
| | | 7 | Mankarso | 0 | 0 | 0 | 0 | 1 | 0 | 0 | 1 | 14.29 (0.4-57.9) |
| | | 11 | Pomona | 0 | 2 | 0 | 0 | 0 | 1 | 0 | 3 | 27.27 (6-61) |
| | | 7 | Pyrogenes | 0 | 0 | 0 | 0 | 1 | 0 | 0 | 1 | 14.29 (0.4-57.9) |
| | | 7 | Wolffi | 1 | 0 | 0 | 0 | 0 | 0 | 0 | 1 | 14.29 (0.4-57.9) |
| **Coyote** | *Canis latrans* | 46 | Australis | 0 | 1 | 0 | 0 | 0 | 0 | 0 | 1 | 2.17 (0.1-11.5) |
| | | 46 | Autumnalis | 3 | 4 | 2 | 1 | 0 | 2 | 0 | 12 | 26.09 (14.3-41.1) |
| | | 74 | Bratislava | 5 | 3 | 1 | 1 | 0 | 0 | 2 | 12 | 16.22 (8.7-26.6) |
| | | 74 | Canicola | 0 | 1 | 0 | 0 | 0 | 0 | 0 | 1 | 1.35 (0-7.3) |
| | | 46 | Cynopteri | 1 | 0 | 0 | 0 | 0 | 0 | 0 | 1 | 2.17 (0.1-11.5) |
| | | 46 | Djasiman | 2 | 3 | 1 | 2 | 0 | 2 | 0 | 10 | 21.74 (10.9-36.4) |
| | | 74 | Grippotyphosa | 2 | 2 | 1 | 0 | 0 | 0 | 0 | 5 | 6.76 (2.2-15.1) |
| | | 38 | Hardjo | 1 | 1 | 0 | 0 | 0 | 0 | 0 | 2 | 5.26 (0.6-17.7) |
| | | 74 | Icterohaemorrhagiae | 4 | 3 | 1 | 1 | 0 | 0 | 1 | 10 | 13.51 (6.7-23.5) |
| | | 46 | Mankarso | 1 | 3 | 1 | 1 | 0 | 0 | 0 | 6 | 13.04 (4.9-26.3) |
| | | 74 | Pomona | 5 | 1 | 3 | 1 | 2 | 0 | 2 | 14 | 18.92 (10.7-29.7) |
| | | 46 | Tarassovi | 1 | 0 | 0 | 0 | 0 | 0 | 0 | 1 | 2.17 (0.1-11.5) |
| **Desert Cottontail** | *Sylvilagus audubonii* | 27 | Georgia | 0 | 0 | 0 | 1 | 0 | 0 | 0 | 1 | 3.7 (0.1-19) |
| | | 30 | Icterohaemorrhagiae | 1 | 0 | 0 | 0 | 0 | 0 | 0 | 1 | 3.33 (0.1-17.2) |
| | | 30 | Pomona | 0 | 1 | 0 | 0 | 0 | 0 | 0 | 1 | 3.33 (0.1-17.2) |
| **Red Fox** | *Vulpes vulpes* | 4 | Autumnalis | 0 | 0 | 1 | 0 | 0 | 0 | 0 | 1 | 25 (0.6-80.6) |
| | | 4 | Djasiman | 0 | 1 | 0 | 0 | 0 | 0 | 0 | 1 | 25 (0.6-80.6) |
| | | 5 | Icterohaemorrhagiae | 1 | 0 | 0 | 0 | 0 | 0 | 0 | 1 | 20 (0.5-71.6) |
| | | 4 | Mankarso | 1 | 0 | 0 | 0 | 0 | 0 | 0 | 1 | 25 (0.6-80.6) |
| | | 5 | Pomona | 0 | 1 | 0 | 0 | 0 | 0 | 0 | 1 | 20 (0.5-71.6) |
| | | 4 | Pyrogenes | 1 | 0 | 0 | 0 | 0 | 0 | 0 | 1 | 25 (0.6-80.6) |
| **Northern Raccoon** | *Procyon lotor* | 16 | Autumnalis | 2 | 0 | 1 | 0 | 0 | 0 | 0 | 3 | 18.75 (4-45.6) |
| | | 134 | Bratislava | 11 | 3 | 9 | 4 | 3 | 3 | 2 | 35 | 26.12 (18.9-34.4) |
| | | 134 | Canicola | 2 | 4 | 2 | 2 | 1 | 0 | 0 | 11 | 8.21 (4.2-14.2) |
| | | 16 | Celledoni | 1 | 0 | 0 | 0 | 0 | 0 | 0 | 1 | 6.25 (0.2-30.2) |
| | | 16 | Cynopteri | 1 | 0 | 0 | 0 | 0 | 0 | 0 | 1 | 6.25 (0.2-30.2) |
| | | 16 | Djasiman | 1 | 1 | 1 | 0 | 0 | 0 | 0 | 3 | 18.75 (4-45.6) |
| | | 134 | Grippotyphosa | 1 | 5 | 4 | 1 | 0 | 0 | 0 | 11 | 8.21 (4.2-14.2) |
| | | 121 | Hardjo | 11 | 5 | 2 | 4 | 3 | 0 | 0 | 25 | 20.66 (13.8-29) |
| | | 134 | Icterohaemorrhagiae | 13 | 9 | 5 | 5 | 1 | 2 | 1 | 36 | 26.87 (19.6-35.2) |
| | | 16 | Mankarso | 0 | 0 | 1 | 0 | 0 | 0 | 0 | 1 | 6.25 (0.2-30.2) |
| | | 134 | Pomona | 6 | 10 | 8 | 9 | 7 | 3 | 11 | 54 | 40.3 (32.2-49.5) |
| | | 16 | Pyrogenes | 1 | 0 | 0 | 0 | 0 | 0 | 0 | 1 | 6.25 (0.2-30.2) |
| | | 16 | Tarassovi | 0 | 0 | 1 | 0 | 0 | 0 | 0 | 1 | 6.25 (0.2-30.2) |
| **Fox Squirrel** | *Sciurus niger* | 69 | Australis | 2 | 0 | 0 | 0 | 0 | 0 | 0 | 2 | 2.9 (0.4-10.1) |
| | | 69 | Autumnalis | 2 | 2 | 0 | 0 | 0 | 0 | 0 | 4 | 5.8 (1.6-14.2) |
| | | 110 | Bratislava | 9 | 15 | 9 | 4 | 2 | 0 | 0 | 39 | 35.45 (26.6-45.1) |
| | | 69 | Celledoni | 1 | 0 | 0 | 0 | 0 | 0 | 0 | 1 | 1.45 (0-7.8) |
| | | 51 | Hardjo | 8 | 16 | 7 | 2 | 1 | 0 | 0 | 34 | 66.67 (52.1-79.2) |
| | | 110 | Icterohaemorrhagiae | 29 | 15 | 9 | 5 | 1 | 1 | 0 | 60 | 54.55 (44.8-64.1) |
| | | 69 | Javanica | 1 | 0 | 0 | 0 | 0 | 0 | 0 | 1 | 1.45 (0-7.8) |
| | | 69 | Mankarso | 1 | 1 | 1 | 0 | 0 | 0 | 0 | 3 | 4.35 (0.9-12.2) |
| | | 110 | Pomona | 1 | 0 | 1 | 0 | 0 | 0 | 0 | 2 | 1.82 (0.2-6.4) |
| | | 69 | Pyrogenes | 2 | 0 | 0 | 0 | 0 | 0 | 0 | 2 | 2.9 (0.4-10.1) |
| **Ground Squirrel** | *Otospermophilus beecheyi* | 4 | Bratislava | 2 | 0 | 0 | 0 | 0 | 0 | 0 | 2 | 50 (6.8-93.2) |
| | | 4 | Hardjo | 0 | 1 | 0 | 0 | 0 | 0 | 0 | 1 | 25 (0.6-80.6) |
| **Striped Skunk** | *Mephitis mephitis* | 8 | Autumnalis | 2 | 0 | 0 | 1 | 0 | 0 | 0 | 3 | 37.5 (8.5-75.5) |
| | | 26 | Bratislava | 0 | 1 | 0 | 0 | 0 | 0 | 0 | 1 | 3.85 (0.1-19.6) |
| | | 8 | Djasiman | 1 | 0 | 1 | 0 | 0 | 0 | 0 | 2 | 25 (3.2-65.1) |
| | | 26 | Icterohaemorrhagiae | 2 | 0 | 1 | 0 | 0 | 0 | 0 | 3 | 11.54 (2.4-30.2) |
| | | 8 | Mankarso | 1 | 0 | 0 | 0 | 0 | 0 | 0 | 1 | 12.5 (0.3-52.7) |
| | | 26 | Pomona | 2 | 0 | 2 | 0 | 1 | 0 | 2 | 7 | 26.92 (11.6-47.8) |
| **Virginia Opossum** | *Didelphis virginiana* | 129 | Bratislava | 1 | 0 | 0 | 0 | 0 | 0 | 0 | 1 | 0.78 (0-4.2) |
| | | 75 | Hardjo | 7 | 0 | 0 | 0 | 0 | 0 | 0 | 7 | 9.33 (3.8-18.3) |
| | | 129 | Icterohaemorrhagiae | 2 | 0 | 0 | 0 | 0 | 0 | 0 | 2 | 1.55 (0.2-5.5) |
| | | 129 | Pomona | 0 | 0 | 0 | 0 | 0 | 0 | 1 | 1 | 0.78 (0-4.2) |

**Table S3: Comparison of *Leptospira* exposure and infection detected in this study and Straub and Foley (2020).** *Leptospira* antibody (MAT) and DNA (PCR) results are shown for our five target species, with lower levels of infection detected in the greater Los Angeles region.

| | Common Name | Scientific Name | ***Leptospira* Exposure (MAT)** | | | ***Leptospira* Infections (PCR)** | | |
|---|---|---|---|---|---|---|---|---|
| | | | ***POS*** | ***n*** | ***%POS (95% CI)*** | ***POS*** | ***n*** | ***%POS (95% CI)*** |
| **This study** | Striped Skunk | *Mephitis mephitis* | 6 | 21 | 28.6 (11.3-52.2) | 5 | 33 | 15.2 (5.1-31.9) |
| | Northern Raccoon | *Procyon lotor* | 31 | 95 | 32.6 (23.4-43.0) | 14 | 162 | 8.6 (4.8-14.1) |
| | Coyote | *Canis latrans* | 14 | 55 | 25.5 (14.7-39.0) | 4 | 107 | 3.7 (1.0-9.3) |
| | Virginia Opossum | *Didelphis virginiana* | 5 | 97 | 5.2 (1.7-11.6) | 1 | 131 | 0.8 (0.0-4.2) |
| | Fox Squirrel | *Sciurus niger* | 66 | 109 | 60.6 (50.7-69.8) | 0 | 148 | 0 (0.0-2.5) |
| **Staub and Foley, 2020** | Striped Skunk | *Mephitis mephitis* | 78 | 206 | 37.9 (31.5-44.7) | 40 | 141 | 28.4 (21.6-36.3) |
| | Northern Raccoon | *Procyon lotor* | 52 | 119 | 43.7 (35.1-52.7) | 23 | 87 | 26.4 (18.3-36.6) |
| | Coyote | *Canis latrans* | 6 | 20 | 30.0 (14.5-51.9) | 2 | 2 | 100.0 (34.2-100.0) |
| | Virginia Opossum | *Didelphis virginiana* | 2 | 32 | 6.3 (1.7-20.1) | 1 | 6 | 16.7 (3.0-56.4) |
| | Fox Squirrel | *Sciurus niger* | 15 | 36 | 41.7 (27.1-57.8) | 4 | 31 | 12.9 (5.1-28.9) |

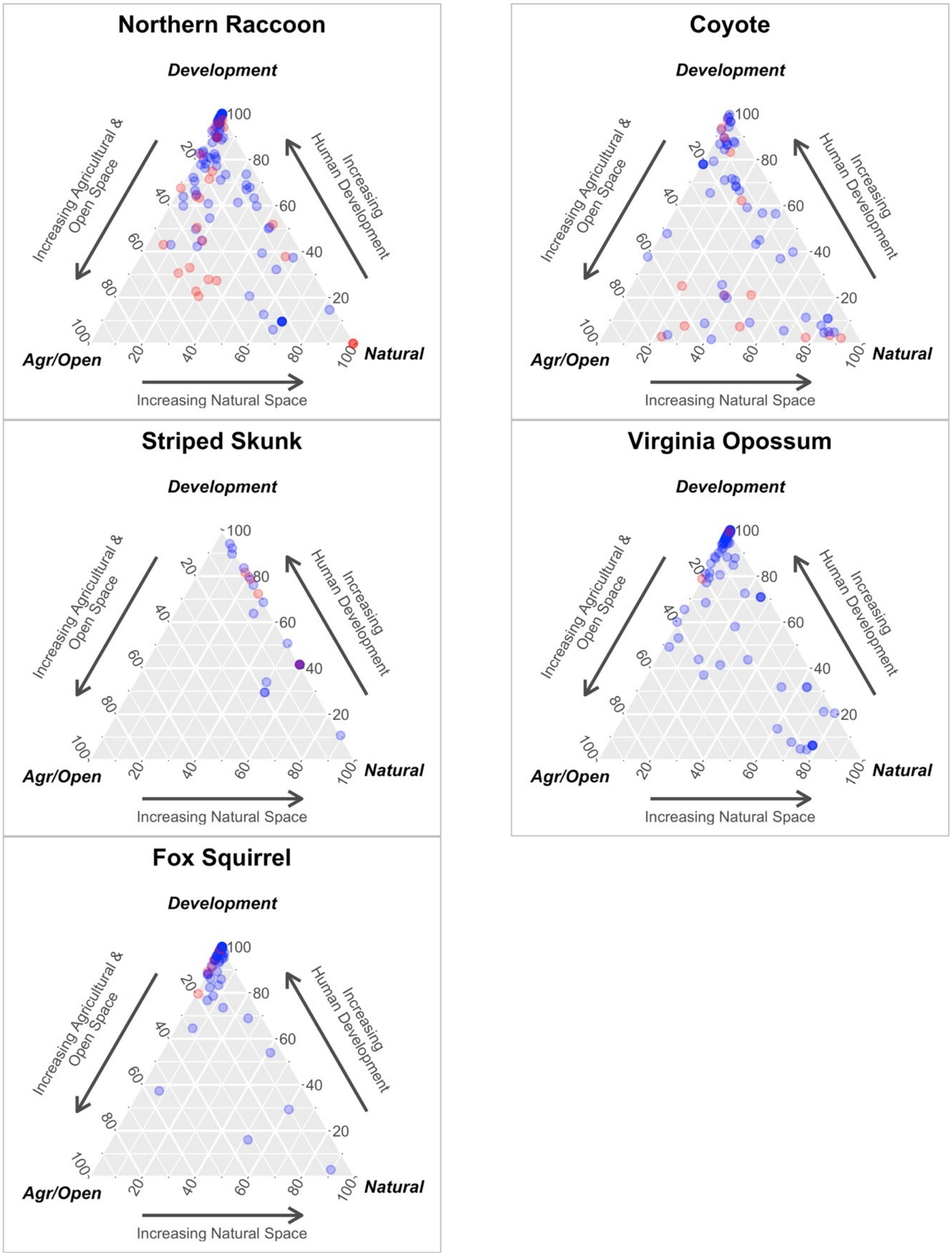


**Figure S1: *Leptospira* antibody results relative to land classification.** Positive (red) and negative (blue) results are shown for each individual species, relative to the proportion of natural, agricultural/open and developed land calculated within the home range buffer of each species.
